# Enhanced mitochondrial fusion during a critical period of synaptic plasticity in adult-born neurons

**DOI:** 10.1101/2023.05.11.540324

**Authors:** Sandra M.V. Kochan, Meret Cepero Malo, Milica Jevtic, Hannah M. Jahn, Gulzar A. Wani, Felix Gaedke, Iris Schäffner, Dieter Chichung-Lie, Astrid Schauss, Matteo Bergami

## Abstract

Integration of new neurons into adult hippocampal circuits is a process coordinated by local and long-range synaptic inputs. To achieve stable integration and uniquely contribute to hippocampal function, immature neurons are endowed with a critical period of heightened synaptic plasticity, yet it remains unclear which mechanisms sustain this form of plasticity during neuronal maturation. We found that, as new neurons enter their critical period, a transient surge in fusion dynamics stabilizes elongated mitochondrial morphologies in dendrites to fuel synaptic plasticity. Conditional ablation of fusion dynamics to prevent mitochondrial elongation selectively impaired spine plasticity and synaptic potentiation, disrupting neuronal competition for stable circuit integration, ultimately leading to decreased survival. Despite profuse mitochondrial fragmentation, manipulation of competition dynamics was sufficient to restore neuronal survival, but left neurons poorly responsive to experiences at the circuit level. Thus, by enabling synaptic plasticity during the critical period, mitochondrial fusion facilitates circuit remodeling by adult-born neurons.

## Introduction

By virtue of their protracted maturation and unique properties, newly-born granule cells (GCs) essentially contribute to circuit information processing in the adult hippocampus (Christian et al., 2014; Miller and Sahay, 2019). Once generated from adult neural stem and progenitor cells (NSPCs), the survival of new GCs is dependent upon competition dynamics throughout their first weeks of life, resulting in the elimination of substantial numbers of neurons and leaving only a fraction of the initially produced GCs stably integrated into the pre-existing hippocampal network (Bergami and Berninger, 2012). At least part of this competition process is believed to be coordinated by activity-dependent secretion of synaptic mediators (neurotransmitters and neurotrophic factors) by pre-existing neurons and, consistent with this notion, local circuit activity and experience can greatly influence the final extent of GCs successfully integrating into the hippocampal network (Gould et al., 1999; Kempermann et al., 1997; van Praag et al., 1999). However, surprisingly little is known about the molecular mechanisms regulating competition between adult-born and pre-existing GCs, or even between cohorts of adult-born GCs of similar age.

An established time window of competition between newborn GCs and the much larger population of pre-existing GCs takes place during the third week of the cell age, when survival is dictated by circuit activity itself (Kleine Borgmann et al., 2016; McAvoy et al., 2016; Tashiro et al., 2006). This is a time characterized by distinctive morphological, ultrastructural and functional traits of plasticity at dendrites and spines of new GCs. It is during this time that the first dendritic spines are formed (Zhao et al., 2006) and new GCs start displaying a greater propensity than their older counterparts for synaptic long-term potentiation (LTP) (Ge et al., 2007; Schmidt-Hieber et al., 2004), a form of synaptic plasticity which underlies memory formation (Whitlock et al., 2006). However, it is only by the fourth week that young GCs transiently engage in multi-synaptic complexes in which several spines get in contact with the same axonal bouton (Toni et al., 2007), a likely structural correlate of ongoing synaptic competition. This time also marks the beginning of a critical period between weeks 4 and 6 of the cell age when LTP reaches its highest magnitude (Ge et al., 2007) and pre-synaptic connectivity remodeling is particularly sensitive to experience (Bergami et al., 2015). While these hallmarks of plasticity are likely to contribute to maintaining the competitiveness of young GCs during their ongoing circuit integration process, which cell-autonomous mechanisms influence this period of enhanced plasticity has remained elusive.

Mitochondria in neurons are known for their roles in regulating energy and metabolite production, Ca^2+^ buffering, reactive oxygen species generation and redox state, all of which are expected to be critical for long-term synapse functioning (Pekkurnaz and Wang, 2022). Mitochondrial structure and function are tightly interwoven (Giacomello et al., 2020), and this is emphasized by the acquisition of compartment-specific differences in the morphological appearance of the mitochondrial network between the axonal and dendritic shafts, with the former exhibiting shorter morphologies and the latter more elaborated ones (Faitg et al., 2021; Popov et al., 2005). In axons, shortening of mitochondria has been linked to their greater transport dynamics from/to synaptic terminals, where they can regulate axonal branching (Courchet et al., 2013; Lewis et al., 2018) and synaptic transmission (Ashrafi et al., 2020; Kang et al., 2008; Kwon et al., 2016; Rangaraju et al., 2014; Sun et al., 2013; Vaccaro et al., 2017). How mitochondria regulate synaptic transmission and plasticity in dendrites is however less understood. Manipulation of mitochondrial content in dendrites appears to be sufficient to elicit changes in spine density (Li et al., 2004; Steib et al., 2014), suggesting a causative effect of mitochondrial function on synaptogenesis. Along these lines, the establishment of larger and stable mitochondrial domains in neurons *in vitro* appears required to sustain protein synthesis during structural plasticity at spines (Rangaraju et al., 2019). Intriguingly, induction of spine plasticity was found to elicit a coincident rapid burst of dendritic mitochondrial fission to facilitate mitochondrial Ca^2+^ uptake and sustain plasticity itself (Divakaruni et al., 2018), implying mitochondrial fission dynamics to play important roles in transducing and coordinating activity-dependent mechanisms at stimulated synapses. Whether other aspects of mitochondrial dynamics may regulate forms of synaptic plasticity, particularly in maturing adult-born GCs *in vivo*, is unknown.

Here, by combining viral-based sensors and genetic approaches to monitor and manipulate mitochondrial dynamics, we report a role for mitochondrial fusion in selectively regulating synaptic plasticity and competitive survival of adult-born GCs during their critical period.

## Results

### A transient period for enhanced mitochondrial fusion in adult-born GCs

To temporally match potential changes in mitochondrial morphology with specific developmental stages during adult-born GC maturation, we took advantage of a retroviral birth-dating approach (Tashiro et al., 2006). We stereotactically injected a combination of two retroviruses encoding for mitochondrial-targeted DsRed (mtDsRed) and cytosolic Turquois (mTurquois2), respectively, followed by confocal imaging and then reconstruction of mitochondria within dendrites of transduced GCs in fixed samples during a time course starting from 2 weeks (i.e., at the beginning of dendritic branching) until 8 weeks of the cell age, when GCs acquired a fully mature phenotype (Figures 1A-1B and S1A). A first qualitative assessment of transduced GCs revealed that while throughout the first 3 weeks of neuronal age dendrites were mostly characterized by shorter and rounder mitochondria that appeared irregularly spaced along the dendritic shaft, during the fourth week mitochondria underwent noticeable changes involving a marked elongation and a more regular distribution along dendrites (Figures 1C and S1A). Consistently, a substantial reduction in the sphericity index of individual mitochondria was detected precisely at 4 weeks of the cell age (Figure 1D). Elongation of mitochondria at 4 weeks also matched a significant increase in mitochondrial occupancy along dendrites (i.e., over 40% of the dendritic volume) as opposed to earlier stages (Figure 1D), indicating that during their fourth week of life adult-born GCs experience a conspicuous remodelling of their dendritic mitochondrial network.

**Figure 1.**
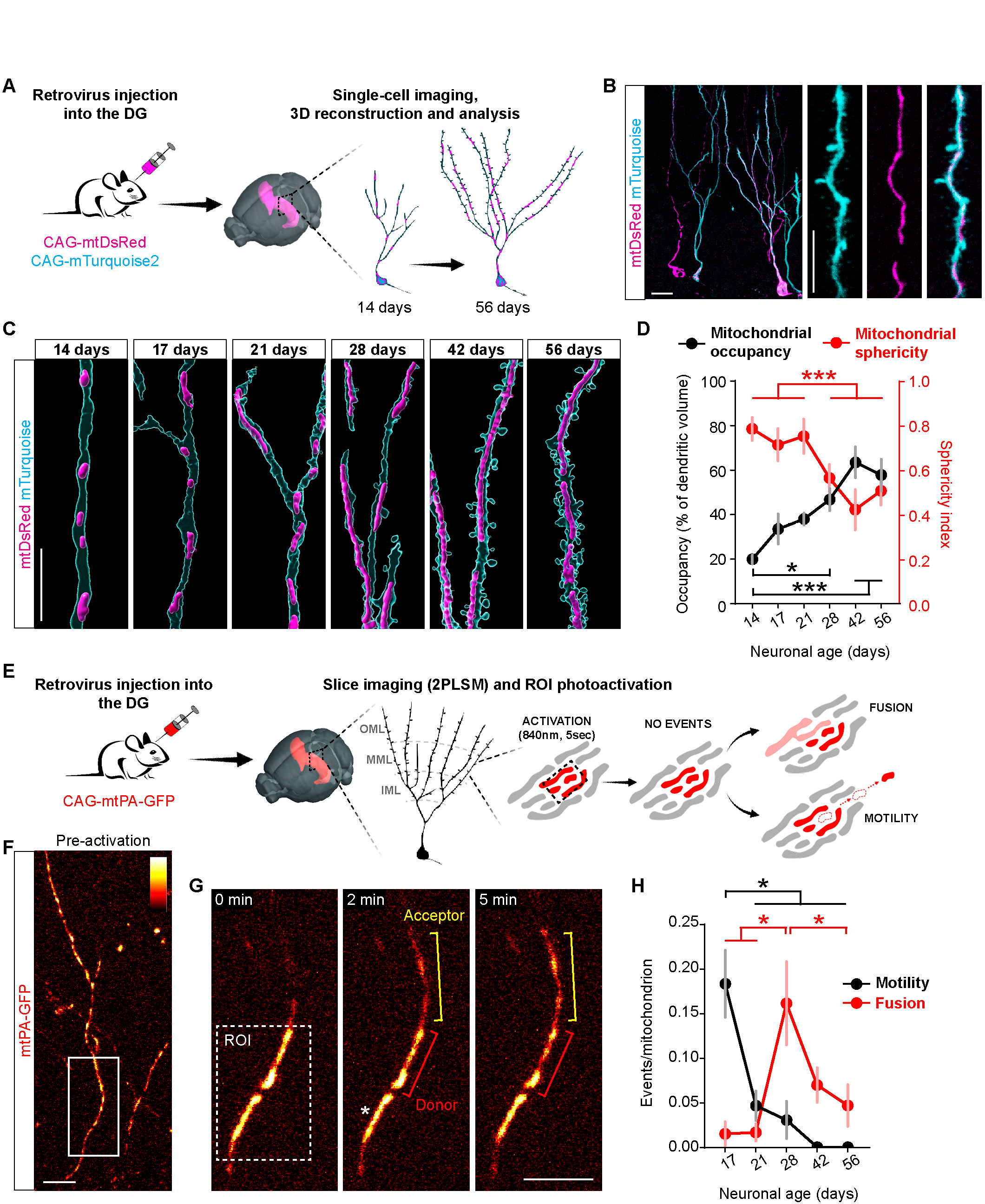
A period for enhanced dendritic mitochondrial fusion in adult-born GCs. **(A)** Experimental setting used for targeting the expression of mtDsRed and mTurquois2 to adult-born GCs via retroviral expression, followed by imaging analysis over the time course of 2 months. **(B)** Example of double-transduced 6-week-old GCs showing mitochondria over dendritic labelling. Bars, 20 and 10 µm. **(C)** Representative images of reconstructed dendritic segments and corresponding mitochondria of adult-born GCs acquired over the course of 2 months. Bar, 10 µm. **(D)** Time course analysis of mitochondrial occupancy in dendrites (percentage of dendritic volume) of adult-born GCs and sphericity index of examined mitochondria from 14 until 56 days of the cell age (n= 10 dendritic segments obtained from 2 mice per time point; one-way Anova followed by Tukey’s multiple comparison test). **(E)** Experimental paradigm used for analyzing mitochondrial dynamics in adult-born GCS via expression of mtPA-GFP and 2PLSM. OML, outer molecular layer; MML, medial molecular layer; IML, inner molecular layer. **(F)** Representative example of a dendritic segment in 28-day-old newborn GC acquired before photoactivation by briefly increasing laser levels at 920 nm. Bar, 10 µm. **(G)** Zoom of the boxed area in F illustrating post-photoactivation of the depicted ROI at time 0 and at following time points. Donor and acceptor mitochondria during a fusion event are indicated. Asterisk refers to a stable mitochondrion undergoing no changes over the examined time window. Bar, 8 µm. **(H)** Time course analysis of mitochondrial motility and fusion proficiency in dendrites of adult-born GCs 17 until 56 days of the cell age (n= 25-48 dendritic ROIs obtained from 2 mice per time point; one-way Anova followed by Tukey’s multiple comparison test). Data are shown as means ± SEM; *, *P* < 0.05; **, *P* < 0.01; ***, *P* < 0.005; See also Figure S1.

Changes in mitochondrial morphology are governed by the concerted action of a set of GTPases (and co-factors) leading to membrane fission and fusion dynamics, whose balance appears to be essential not only in maintaining mitochondrial function but also in adjusting mitochondrial size to meet cellular and metabolic demands (Giacomello et al., 2020). To systematically investigate the extent of mitochondrial dynamics in maturing adult-born neurons in real-time, we engineered a retrovirus to encode a photo-activatable green fluorescent protein (mtPA-GFP) targeted to the mitochondrial matrix (Karbowski et al., 2004). At the desired time points after retroviral injection, acute hippocampal slices were prepared and transduced GCs identified utilizing 2-photon laser-scanning microscopy (2PLSM) (Figure 1E). Visual analysis of the whole mitochondrial network in GCs expressing mtPA-GFP confirmed a cell age-dependent transformation of the mitochondrial network morphology in neuronal dendrites (Figure S1B-S1C). Photo-activation was then carried out in specific regions of interests (ROIs) along the dendritic arbor of transduced GCs, typically in portions of the branches located within the medial molecular layer (MML) (Figure 1E). This allowed us to track and analyse photo-activated mitochondria whose signal intensity specifically and markedly increased, and could therefore be easily distinguished from non-photo-activated adjacent organelles (Göbel et al., 2020) (Figure 1E). Subsequent time-lapse imaging of photo-activated ROIs over the course of 20 minutes allowed us to recognize two distinctive groups of events in mitochondrial behaviour: fusion and motility events. Moving mitochondria were observed as photo-activated organelles that would travel within (and/or leave) the initially photo-activated dendritic ROI, without losing signal intensity. These were typically small mitochondria that significantly migrated (> 5μm in distance) either retrogradely or anterogradely along the dendritic shaft, and could be distinguished by kymograph analysis (Figure S1D). Interestingly, and in line with previous studies in other neuronal types (Faits et al., 2016), these events were readily detectable at early time points but their frequency rapidly declined with the maturation of neurons to become virtually undetectable by 4-6 weeks (Figure 1H **and S1D**), disclosing a progressive stabilization of dendritic mitochondria as these became more elongated. In net contrast, photo-activated mitochondria (which we termed “donors”) undergoing fusion events did not typically move, were larger in size and showed a quantifiable decline in GFP intensity which was temporally matched by an opposite increase in brightness of nearby fusing mitochondria, which had not been initially photo-activated (termed “acceptors”) (Figure 1F-1G and S1D-S1E). Intriguingly, while GCs of 3 weeks of age or younger, as well as GCs older than 6 weeks, only rarely exhibited fusion events during the imaging session, the fourth week of age marked a period of net increase in the frequency of mitochondrial fusion (Figure 1H).

Taken together, these results indicate that adult-born GCs transiently experience heightened rates of mitochondrial fusion during their fourth week of age, shortly before exhibiting an elongation and stabilization of their dendritic mitochondria.

### Manipulation of mitochondrial fusion dynamics disrupts dendritic mitochondrial domains

To causally link developmental changes in mitochondrial fusion dynamics with the formation of stable, larger mitochondrial domains in the dendrites of adult-born GCs, we conditionally manipulated the genes responsible for outer mitochondrial membrane fusion *Mitofusin1* (*Mfn1*) and *Mfn2* utilizing established mouse lines carrying a floxed exon in each of these alleles (Lee et al., 2012). We opted for a Cre and mtDsRed-expressing retrovirus-based approach to obtain a sparse, single-cell conditional knock-out (cKO) for *Mfn1* or *Mfn2*, and then monitored changes in the mitochondrial network of transduced GCs *in vivo* (Figure 2A). By 6 weeks after stereotactic injection, when mitochondria had achieved stable and elongated morphologies in control GCs (Figures 1C and 1H), Mfn1^cKO^ and Mfn2^cKO^ GCs showed a marked network fragmentation throughout their dendritic tree and cell body (Figure 2B). Despite being fragmented, mitochondria did not exhibit overtly enlarged or swollen morphologies which are characteristic of progressive mitochondrial dysfunction, as previously described in other *in vivo* neuronal systems following chronic ablation of mitochondrial fusion (Chen et al., 2007; Lee et al., 2012; Motori et al., 2020), thus suggesting the 6-week time point in our settings to mark a rather early stage of disrupted organelle morphology. Yet, analysis of dendrites in mutant GCs revealed a greater depletion of mitochondria with increasing distance from the cell soma, that is, at more distally located dendritic regions (Figures 2C and 2D). Consistently, more proximal tracts of the dendritic arbor located in the inner molecular layer (IML) appeared virtually undistinguishable from control neurons in terms of mitochondrial occupancy, whereas more distal dendritic portions in the MML and outer molecular layer (OML) were particularly affected, leaving significant segments of dendrites seemingly devoid of mitochondria (Figures 2D and 2E). Intriguingly, simultaneous deletion of both *Mfn1* and *Mfn2* did not result in an overtly worsened phenotype as compared to single mutants (Figure 2B-2E).

**Figure 2.**
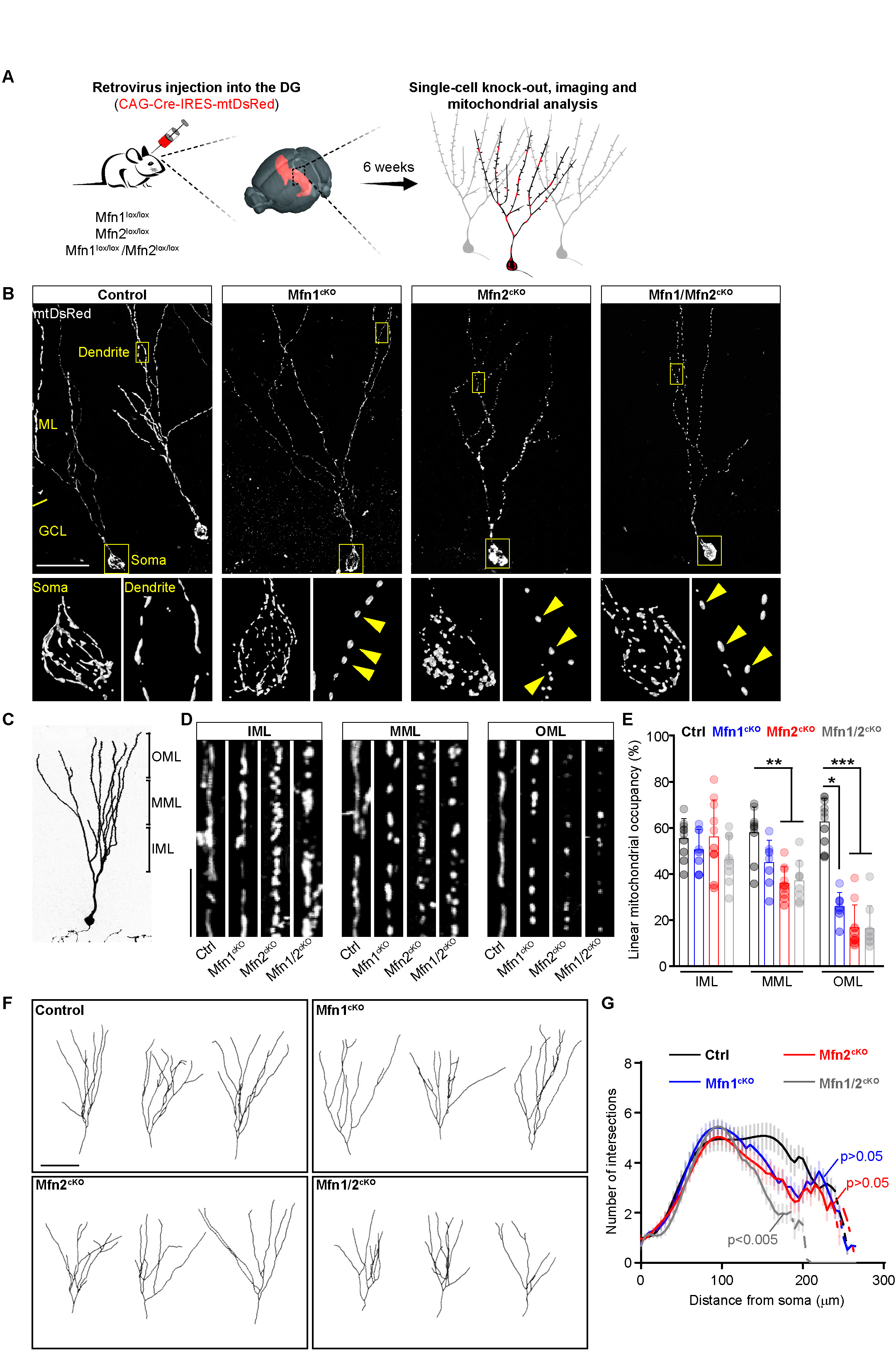
Disruption of dendritic mitochondrial domains by manipulation of *Mfn1* and *Mfn2*. **(A)** Experimental setting used for simultaneous retroviral expression of Cre and mtDsRed in Mfn1^lox/lox^, Mfn2 ^lox/lox^ and Mfn1^lox/lox^ /Mfn2^lox/lox^ mice and subsequent analysis of mitochondrial network morphology at 6 weeks. **(B)** Examples of transduced GCs showing changes in the mitochondrial network following gene deletion. Lower panels depict zooms of the boxed area for soma and dendrite. Arrowheads point to fragmented mitochondria along dendrites. Bar, 50 µm. ML, molecular layer; GCL, granule cell layer. **(C)** Picture of a GC illustrating location of dendrites according to layers in the ML. OML, outer molecular layer; MML, medial molecular layer; IML, inner molecular layer. **(D)** Magnification of transduced dendritic segments located in the three layers of the ML for Mfn1^cKO^, Mfn2^cKO^ and Mfn1/2^cKO^ GCs. Bar, 10 µm. **(E)** Quantification of linear dendritic mitochondrial occupancy in transduced Mfn1^cKO^, Mfn2^cKO^ and Mfn1/2^cKO^ GCs across layers of the ML (n= 8-10 neurons obtained from 2 mice per time point; Kruskal-Wallis test followed by Dunn’s multiple comparison test). **(F)** Representative reconstructions of the dendritic arbor of GCs transduced with GFP-Cre and DsRed for each of the indicated genotypes. Bar, 50 µm. **(G)** Sholl analysis of transduced Mfn1^cKO^, Mfn2^cKO^ and Mfn1/2^cKO^ GCs illustrating dendritic arbor complexity (n= 15 neurons obtained from 3-5 mice per genotype; Kolmogorov-Smirnov test). Data are shown as means ± SEM; *, *P* < 0.05; **, *P* < 0.01; ***, *P* < 0.005; See also Figure S2.

To understand if the introduced genetic manipulations may interfere with the morphological development of dendrites, we then utilized a combination of cytosolic DsRed and nuclear Cre-GFP-encoding retroviruses to reconstruct the dendritic tree of wild-type versus knock-out neurons double-labelled for DsRed and GFP for each of the examined mouse lines at 6 weeks after viral injection (Figure 2F). Although the occurrence of double-transduced (DsRed+ and Cre-GFP+) Mfn1^cKO^, Mfn2^cKO^ and Mfn1/2^cKO^ GCs was visibly reduced as compared to that of wild-type mice, analysis showed that the number of dendritic branches and branching points was mostly unaffected between genotypes (Figure S2A-S2C). In contrast, Mfn1/2^cKO^ GCs exhibited a significant shortening of their total dendritic length, suggesting that complete ablation of mitochondrial fusion may interfere with the growth of more distal dendritic branches (Figure S2A-S2C). In line with these quantifications, Sholl analysis disclosed mild changes in the overall dendritic complexity of Mfn1^cKO^ or Mfn2^cKO^ neurons, while double mutant GCs exhibited a significant impairment, particularly in most distal regions of the dendritic tree (Figures 2F and 2G).

Thus, in contrast to Mfn1/2^cKO^ GCs, disruption of mitochondrial fusion dynamics in adult-born GCs via deletion of either *Mfn1* or *Mfn2* is sufficient to prevent the formation of elongated dendritic mitochondrial domains without overly impairing the genesis, overall complexity or integrity of dendrites.

### Loss of mitochondrial fusion impairs adult-born GC incorporation into the pre-existing network

The apparent reduced frequency at which virally transduced Mfn1^cKO^, Mfn2^cKO^ and Mfn1/2^cKO^ GCs were found in our samples during the analysis of dendritic complexity prompted us to investigate potential defects in GC survival. To this aim, we quantified the total amount of transduced GCs in *Mfn1^lox/lox^*, *Mfn2^lox/lox^* or *Mfn1/Mfn2^lox/lox^* mice following combined injection of fixed volumes of CAG-GFP-Cre (fusion protein) and CAG-DsRed-encoding retroviruses of known titre (Figures 3A and 3B). Normalization of the relative proportions of co-transduced (knockout) new neurons (GFP-Cre+/DsRed+) onto all DsRed+ neurons allowed us to calculate a survival index regardless of potential small variations in injection site and titre between used viruses (Jagasia et al., 2009; Tashiro et al., 2006) (Figure 3A). Analysis of these samples revealed no alterations in the survival index for any of the examined genotypes at 3 weeks of the GC age (Figure 3C), a time when the mitochondrial network is still largely fragmented also in wild-type GCs (Figures 1C and S1A). However, by 6 weeks the survival ratio of Mfn1^cKO^, Mfn2^cKO^ and Mfn1/2^cKO^ GCs was reduced by about 50% as compared to transduced GCs in control mice (Figures 3B and 3C). Of note, the remaining surviving fraction of mutant GCs (GFP-Cre+/DsRed+) appeared unaffected in terms of neurite integrity in all examined genotypes, and exhibited no evident signs of degeneration (Figure 3D). Lastly, specificity of this survival effect was confirmed in *Mfn2^lox/lox^* mice by replacing the CAG-GFP-Cre virus with a virus co-expressing *Mfn2* cDNA and GFP-Cre (CAG-Mfn2-T2A-GFP-Cre, referred to as Mfn2 OE), which by 6 weeks was sufficient to restore GC survival to levels comparable to 3 week-old Mfn2^cKO^ GCs and to that of control mice (Figure 3C), without altering GC morphology and positioning in the GCL (Figure S2D).

**Figure 3.**
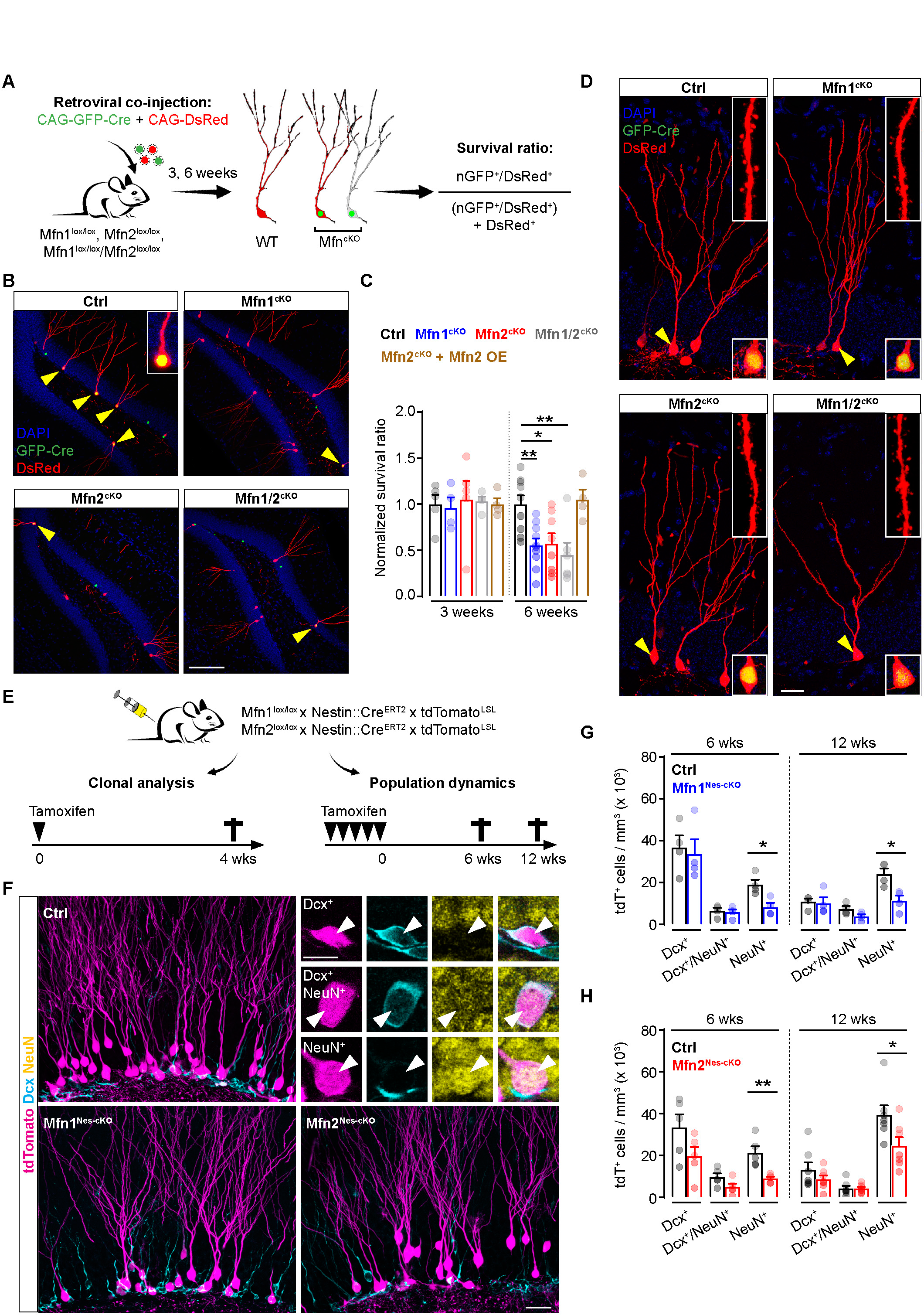
Loss of mitochondrial fusion impairs adult-born GC circuit incorporation. **(A)** Experimental setting used for simultaneous injection of CAG-GFP-Cre and CAG-DsRed retroviruses in Mfn1^lox/lox^, Mfn2 ^lox/lox^ and Mfn1^lox/lox^ /Mfn2^lox/lox^ mice and subsequent survival index analysis at 3 and 6 weeks. **(B)** Pictures of the DG illustrating examples of co-transduction rates in the indicated genotypes at 6 weeks. Arrowheads and inset point to double-transduced GCs. Bar, 100 µm. **(C)** Quantification of survival index following co-injection of CAG-DsRed and CAG-GFP-Cre (or CAG-Mfn2-T2A-GFP-Cre, for simplicity referred to as Mfn2 OE) retroviruses in the indicated genotypes at 3 and 6 weeks (n= 4-10 mice per condition; one-way Anova followed by Tukey’s multiple comparison test). **(D)** Representative pictures of transduced GCs expressing GFP-Cre and DsRed in the indicated genotypes at 6 weeks. Inset shows a zoom of a dendritic segment in the MML. Arrowheads point to double-transduced GCs among other DsRed-only positive GCs. Bar, 20 µm. **(E)** Experimental paradigms used for clonal analysis and assessment of population dynamics in Mfn1^Nes-cKO^ and Mfn2^Nes-cKO^. **(F)** Representative pictures of the superior DG blade showing tdTomato reporter+ GCs superimposed to Dcx immunostaining in the indicated genotypes. Top right panels depict examples of tdTomato+ newborn GCs being immunoreactive for Dcx, NeuN or both. Bars, 25 and 10 µm. **(G-H)** Quantification of tdTomato+ GCs expressing Dcx, NeuN or both in Mfn1^Nes-cKO^ and Mfn2^Nes-cKO^ mice at 6 and 12 weeks after tamoxifen treatment (n= 4-7 mice per condition and time point; Mann-Whitney test). Data are shown as means ± SEM; *, *P* < 0.05; **, *P* < 0.01; See also Figures S2 and S3.

We next sought to independently validate these findings by generating tamoxifen-inducible conditional knock-out mouse lines for *Mfn1* (termed Mfn1^Nes-cKO^) and *Mfn2* (Mfn2^Nes-cKO^), in which Cre^ERT2^ expression is controlled by the Nestin promoter (Nestin::Cre^ERT2^) (Lagace et al., 2007), and whose recombination is revealed by a cytosolic tdTomato reporter (Madisen et al., 2010) (Figure 3E). As the initial recombination in these mice takes place in NSPCs, and is then retained life-long in their neuronal progeny (i.e., GCs), we first examined potential changes in adult NSPC activity resulting from tamoxifen-induced ablation of *Mfn1* or *Mfn2* that may in turn affect the production of new GCs. Assessment of NSPC behaviour was conducted by treating these animals with a single low-concentration dose of tamoxifen sufficient to induce sparse labelling of radial glia-like NSPCs (which we termed “R”), allowing us to fate-map their activity at the single clone level over the course of 1 month (Bonaguidi et al., 2011) (Figures 3E and S2E). Analysis of sections from these treated mice disclosed no differences between genotypes in the proportions of clones exhibiting “quiescent” (clones containing isolated Rs, which did not divide over the examined period of time), “active” (clones in which Rs underwent one or multiple rounds of division, including self-renewal) or “depleted” (clones in which Rs were active but became ultimately depleted from the clone) phenotypes (Figures S2E and S2F). Likewise, the proportion of “active” clones involving exclusively self-renewing (clones containing only Rs), neurogenic (Rs plus neurons), gliogenic (Rs plus astrocytes) or mixed (neurogenic and gliogenic) behaviours (Figure S2G), as well as the average clone size (Figure S2H), were virtually unchanged in Mfn1^Nes-cKO^ and Mfn2^Nes-cKO^ mice as compared to controls.

Importantly, we corroborated these data by taking advantage of two additional Cre-expressing mouse lines targeting adult NSPCs. In the first of these, Cre^ER^ expression is controlled by the hGFAP promoter (hGFAP::Cre^ER^) (Chow et al., 2008), which is active in NSPCs (Wani et al., 2022), and recombination was revealed by crossing this line with a mtYFP reporter mouse (*mtYFP^LSL^* mice) (Sterky et al., 2011) to generate Mfn1^GFAP-cKO^ and Mfn2^GFAP-cKO^ mice (Figure S2I). In these mice, morphologically identified radial GFAP+ NSPCs whose soma was located in the sub-granular zone (SGZ) of the DG exhibited widespread mitochondrial fragmentation following ablation of either *Mfn1* or *Mfn2* (Figure S2J). Yet, BrdU incorporation analysis conducted at 4 weeks after tamoxifen treatment (Figure S2I) revealed that NSPC proliferation was not affected in Mfn1^GFAP-cKO^ and Mfn2^GFAP-cKO^ mice (Figures S2K and S2L), despite pronounced mitochondrial fragmentation. In contrast, simultaneous deletion of both, *Mfn1* and *Mfn2* (Mfn1/2^GFAP-cKO^) caused a significant reduction of NSPC proliferation (Figures S2K and S2L), in line with previous studies (Khacho et al., 2016), thus suggesting that ablation of both genes is required to significantly alter NSPC fate. Lastly, lack of specific effects in adult NSPCs subjected to *Mfn2* ablation was verified by using Mfn2^Glast-cKO^ mice, which were generated by using the inducible Cre driver line Glast::Cre^ERT2^ mice (Mori et al., 2006). Consistent with our prior analysis in Mfn2^Nes-cKO^ mice, repetitive tamoxifen treatment did not alter the overall density of recombined radial glia-like NSPCs co-expressing the tdTomato genetic reporter for Cre activity and the endogenous NSPC marker Sox2 up to 12 weeks after tamoxifen treatment (Figures S3A and S3B). We conclude that, taken individually, *Mfn1* and *Mfn2* are largely dispensable for adult NSPC activity and the early proliferative steps of neurogenesis, thus establishing the above-mentioned lines as suitable models for investigating the specific role of mitochondrial fusion in the development and survival of adult-born GCs.

Repetitive tamoxifen treatment of Mfn1^Nes-cKO^ and Mfn2^Nes-cKO^ mice did not alter the pool of tdTomato+ radial glia-like NSPCs (Figures S3C and S3D), allowing us to systematically examine potential changes in population dynamics of newly-generated GCs (Figure 3E). Confocal analysis of brain sections obtained from these mice revealed a discernible reduction in the number of tdTomato+ GCs in both, Mfn1^Nes-cKO^ and Mfn2^Nes-cKO^ mice, despite absence of any overt signs of degeneration in the surviving neurons (Figure 3F), which is consistent with the data collected from viral injections (Figures 3D). To investigate potential alterations in the maturation process of tdTomato+ GCs, we examined the expression of cellular markers such as doublecortin (Dcx), which is expressed in young immature GCs up to the fourth week of age (Brown et al., 2003), and neuronal nuclear protein (NeuN), which starts to be expressed as Dcx declines during the acquisition of a more mature phenotype (Figure 3F). In line with our viral experiments conducted at 3 weeks of the GC age (Figure 3C), the pool of immature Dcx+ (and Dcx/NeuN co-expressing) GCs did not appear significantly affected by *Mfn1* or *Mfn2* ablation at 6 and 12 weeks after tamoxifen-induced recombination (Figure 3G and 3H). This is consistent with the comparable rates of neurogenesis exhibited by Mfn1^Nes-cKO^, Mfn2^Nes-cKO^ and control littermates (Figures S2F and S2G). In contrast, quantification of tdTomato+ GCs that had fully become NeuN+ in mutant mice revealed that their density was essentially halved as compared to control mice at both examined time points (Figures 3G and 3H). In line with this finding, TUNEL staining disclosed an evident (yet not significant) increase in the number of tdTomato+ cells undergoing cell death as compared to controls (Figure S3E).

Together, these data demonstrate that while neurogenesis and the early maturation steps of new GCs are not particularly affected by *Mfn1* or *Mfn2* ablation, by the time fusion-deficient GCs exit the fourth week of age their incorporation into the pre-existing network becomes compromised.

### Defective synaptic plasticity in GCs lacking mitochondrial fusion

The specific timing at which the reduction in GC numbers became manifest (Figures 3C, 3G and 3H), together with the absence of explicit hallmarks of neurodegeneration in all of the examined models lacking mitochondrial fusion led us to consider alternative possibilities that may explain the negative selection of mutant GCs. As the timing of reduced GC survival largely overlapped with the critical period of post-synaptic plasticity that occurs between 4 and 6 weeks of GC age (Ge et al., 2007), we reasoned that changes in GC viability may therefore result from a primary impairment in synaptic integration and/or plasticity rather than severe mitochondrial dysfunction, which would otherwise be expected to cause profuse neurodegeneration (Chen et al., 2007; Motori et al., 2020). Consistent with this notion, the first discernible signs of degeneration (i.e., neurite beading, appearance of apoptotic cell bodies and condensed/pyknotic nuclei) in adult-born GCs lacking *Mfn2* only appeared around 6 months after tamoxifen treatment (Figure S3F), that is, over 4 months after the first significant drop in GC numbers, arguing for a distinct mechanism regulating the survival of adult-born GCs lacking mitochondrial fusion at 6 weeks of age.

To gain insights into the potential consequences of disrupted mitochondrial fusion for the synaptic integration of GCs, we first examined dendritic spines. The overall spine density in the MML of Mfn1^Nes-cKO^ and Mfn2^Nes-cKO^ mice appeared undistinguishable from that of controls up to 12 weeks after tamoxifen treatment (Figures 4A and 4B), indicating that spine formation can occur in absence of mitochondrial fusion. Likewise, correlative light and electron microscopy (CLEM) analysis of randomly chosen tdTomato+ dendritic segments in Mfn2^Nes-cKO^ mice (Figure S3G) showed a seemingly normal ultrastructure of spines and synapses (Figures S3H). Yet, GCs lacking *Mfn1* or *Mfn2* exhibited a net reduction in the occurrence of larger “mushroom” spines (i.e., spines with an area equal or greater than 0.4 μm^2^) (Zhao et al., 2006) (Figure 4A and 4C), which are generally regarded as a hallmark of structural synaptic plasticity during long-term potentiation (LTP) (Matsuzaki et al., 2004). Focusing on *Mfn2*, we obtained similar results by analysing 6-week-old Mfn2^cKO^ GCs that had been previously birth-dated by viral injections. Specifically, Mfn2^cKO^ GCs failed in up-regulating mushroom spine density when mice were exposed for 4 weeks to environmental enrichment (EE, a condition known to promote synaptic plasticity in adult-born GCs) (Bergami et al., 2015; Zhao et al., 2014) (Figure 4D), in contrast to GCs of wild-type mice, in which mushroom spine density almost doubled during this period (Figures 4E and 4F). Conversely, to test if enhancing mitochondrial fusion would be sufficient to elicit spine plasticity, we retrovirally overexpressed *Mfn2* in wild-type mice maintained in standard housing. *Mfn2* overexpression did not result in an abnormally hyperfused mitochondrial network, yet transduced GCs showed a noticeable elongation of dendritic mitochondria by 3 weeks of the cell age (Figure S3I), a time when control GCs still exhibited a largely fragmented network (Figure S1A and S3I). By this age, *Mfn2* overexpression was sufficient to significantly increase mushroom spine density as compared to control GCs, which at 3 weeks are very immature and only exhibit very few mushroom spines (Zhao et al., 2006) (Figure 4G **and S3J**). However, prolonged *Mfn2*-overexpression failed to elicit a further increase in mushroom spine density in 6-week-old GCs (Figure 4G), a time when maturation is almost complete and mitochondria have achieved stable and elaborated morphologies also in control GCs (Figure 1C **and S1A**), suggesting that forcing mitochondrial fusion is sufficient to accelerate spine maturation and plasticity only during GC development. Thus, while mitochondrial fusion is dispensable for dendritic integrity and attaining stable spine density, it is necessary for spine plasticity in adult-born GCs.

**Figure 4.**
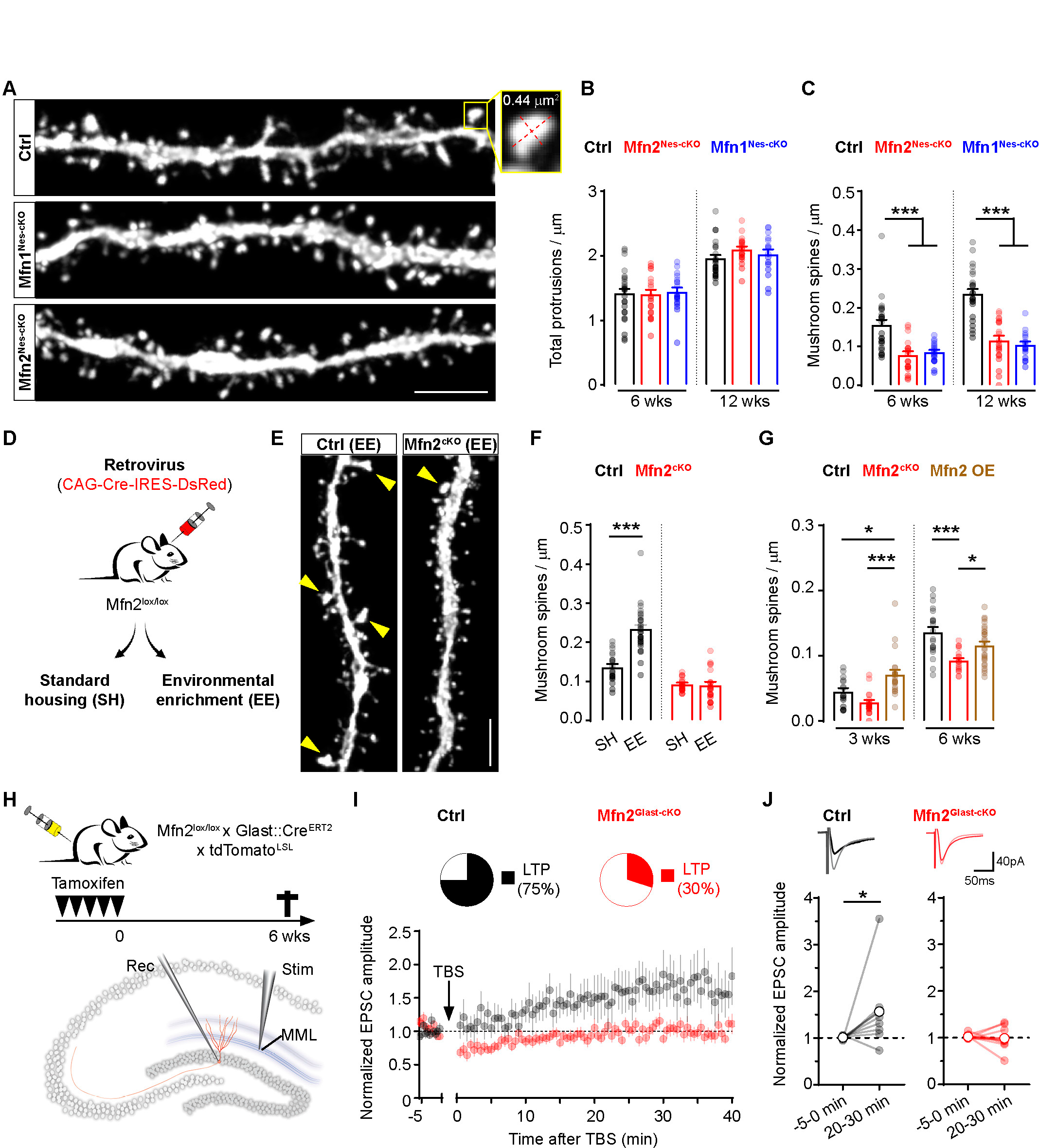
Defective synaptic plasticity in GCs lacking mitochondrial fusion. **(A)** Examples of dendritic spines in the MML of tdTomato+ GCs in Mfn1^Nes-cKO^, Mfn2^Nes-cKO^ and control littermates at 6 weeks. Zoom on the right shows a mushroom spine, defined as having a spine head area equal or greater than 0.4 μm^2^. Bar, 5 µm. **(B-C)** Quantification of total dendritic protrusions and mushroom spines in Mfn1^Nes-cKO^ and Mfn2^Nes-cKO^ mice at 6 and 12 weeks after tamoxifen treatment (n= 16-27 dendritic segments obtained from 3-5 mice per condition and time point; Kruskal-Wallis test followed by Dunn’s multiple comparison test). **(D)** Scheme showing the paradigm used for examining spine density in Mfn2^lox/lox^ mice exposed to environmental enrichment (EE) via retroviral co-expression of Cre and cytosolic DsRed. **(E)** Examples of dendritic spines in the MML of DsRed+ Mfn2^cKO^ and control GCs after 4 weeks of EE. Arrowheads point to mushroom spines. Bar, 5 µm. **(F)** Quantification of mushroom spine density in GCs of Mfn2^cKO^ and control mice maintained in standard housing (SH) or exposed to EE for 4 weeks (n= 18-28 dendritic segments obtained from 3-4 mice per condition and time point; Mann-Whitney test). **(G)** Quantification of mushroom spine density in GCs of Mfn2^cKO^ and control mice transduced with CAG-Cre-IRES-DsRed and maintained in standard housing (SH) or exposed to EE for 4 weeks. Co-injection of CAG-Mfn2-T2A-GFP (referred to as Mfn2 OE for simplicity) was used to over-express Mfn2 in control mice (n= 15-26 dendritic segments obtained from 3 mice per condition and time point; Kruskal-Wallis test followed by Dunn’s multiple comparison test). **(H)** Scheme showing the paradigm used for whole-cell patch-clamp recordings of passive and active membrane properties of GCs as well as for induction of synaptic plasticity following stimulation of the MML in the DG of acute hippocampal slices obtained from Mfn2^Glast-cKO^ and control mice at 6 weeks after tamoxifen treatment. **(I)** Quantification of averaged EPSC amplitudes recorded in whole-cell voltage-clamp from tdTomato+ GCs following LTP induction by theta-burst stimulation (TBS, arrow at time 0), and normalized to baseline recordings obtained before LTP induction. Upper pie charts depicts the percentage of GCs that exhibited LTP after induction for Mfn2^cKO^ (n= 10 neurons recorded from 9 mice) and control mice (n= 8 neurons recorded from 7 mice; Kruskal-Wallis test followed by Dunn’s multiple comparison test). **(J)** Plot depicting individual (light color) and mean EPSCs potentiation of recorded GCs before and after (time 20-30 min during recording) TBS (n= 8-10 neurons recorded from 7-9 mice; Wilcoxon test). Top traces show examples of baseline ESPCs (solid color) versus LTP (light color) of EPSCs in control and Mfn2^cKO^ GCs. Data are shown as means ± SEM; *, *P* < 0.05; ***, *P* < 0.005; See also Figures S3 and S4.

To functionally corroborate these structural data, we conducted whole-cell patch-clamp recordings of adult-born GCs lacking *Mfn2* (Mfn2^Glast-cKO^ mice at 6 weeks post-tamoxifen treatment) and assessed their electrophysiological and synaptic properties in acute brain slices (Figure 4H). Analysis revealed that passive membrane properties (Figure S4A), including resting membrane potential (E_M_), input resistance (R_input_) and membrane time constant (τ_m_), which are good indicators of GC maturation, as well as threshold and amplitude of action potentials (AP) (Figure S4B) and current-voltage relationship (Figure S4C) were undistinguishable between mutant and control GCs. GCs defective for *Mfn2* were altogether less excitable than controls (Figure S4D), yet they exhibited a significantly higher frequency of spontaneous inputs (sPSCs), albeit of lower average amplitude (Figure S4E). We next monitored potential changes in synaptic plasticity of new GCs, and first examined their response to paired-pulse stimulation of afferent fibres (perforant path) located in the MML (Figure 4H), a paradigm utilized to induce a form of short-term presynaptic plasticity (Zucker and Regehr, 2002). We observed a modest facilitation of recorded excitatory post-synaptic currents (EPSCs), which was however similar between groups (Figure S4F). Then, we examined synaptic responses following application of a theta-burst stimulation (TBS) to the MML, a widely used paradigm to induce synaptic LTP in adult-born GCs (Ge et al., 2007; Schmidt-Hieber et al., 2004). Intriguingly, while TBS induced a potentiation of baseline EPSCs lasting for the duration of the recording (about 40-45 minutes) in about 75% of control GCs, only 30% of *Mfn2*-deficient neurons underwent LTP and, when present, LTP exhibited lower amplitudes (Figure 4I and 4J), disclosing defects in the induction of postsynaptic plasticity.

Taken together, these results support the notion that mitochondrial fusion is required for activity- and experience-dependent forms of synaptic plasticity in adult-born GCs.

### Mitochondrial fusion regulates competition dynamics between adult-born GCs

As competition between cohorts of GCs is regulated at the synaptic level (Kleine Borgmann et al., 2016; McAvoy et al., 2016; Tashiro et al., 2006), the transiently heightened synaptic plasticity of newborn GCs has been suggested to contribute to enabling their integration into the pre-existing circuits (Ge et al., 2007; Toni et al., 2007). To understand whether the enhanced mitochondrial fusion rates at the beginning of the critical period of plasticity may contribute to competition dynamics, we experimentally manipulated the extent of new GCs lacking *Mfn2* by two independent approaches. First, by varying tamoxifen dosages (full dosage of 5 injections over 5 consecutive days versus a single tamoxifen injection) administered to Mfn2^Glast-cKO^ mice we were able to achieve either nearly complete (over 95%) or only partial (about 56%) recombination in new GCs (as assessed by tdTomato expression), with the latter condition leaving a significant fraction of new GCs being not recombined (i.e., tdTomato-negative, mitochondrial fusion-proficient GCs) (Figure 5A and 5B). Quantification of Dcx+ and NeuN+ GCs under these two conditions disclosed that partial recombination led to a significant drop (54%) in the density of tdTomato+/NeuN+ GCs lacking *Mfn2* (Figure 5C and 5D), mirroring the phenotype previously observed in Mfn2^Nes-cKO^ mice (Figure 3H), in which maximal penetrance of Cre-mediated recombination even following full-dosage tamoxifen treatment did not exceed 40% (Figure 5A). In stark contrast, a virtually complete recombination in Mfn2^Glast-cKO^ mice was sufficient to restore the density of tdTomato+/NeuN+ GCs to levels indistinguishable from those of control mice (Figure 5C and 5D). These results confirm that the survival effects caused by *Mfn2* deletion at the time point of 6 weeks are not due to neurodegeneration, and rather suggest a role for the relative expression of MFN2 between GC of similar ages in regulating competitive survival.

**Figure 5.**
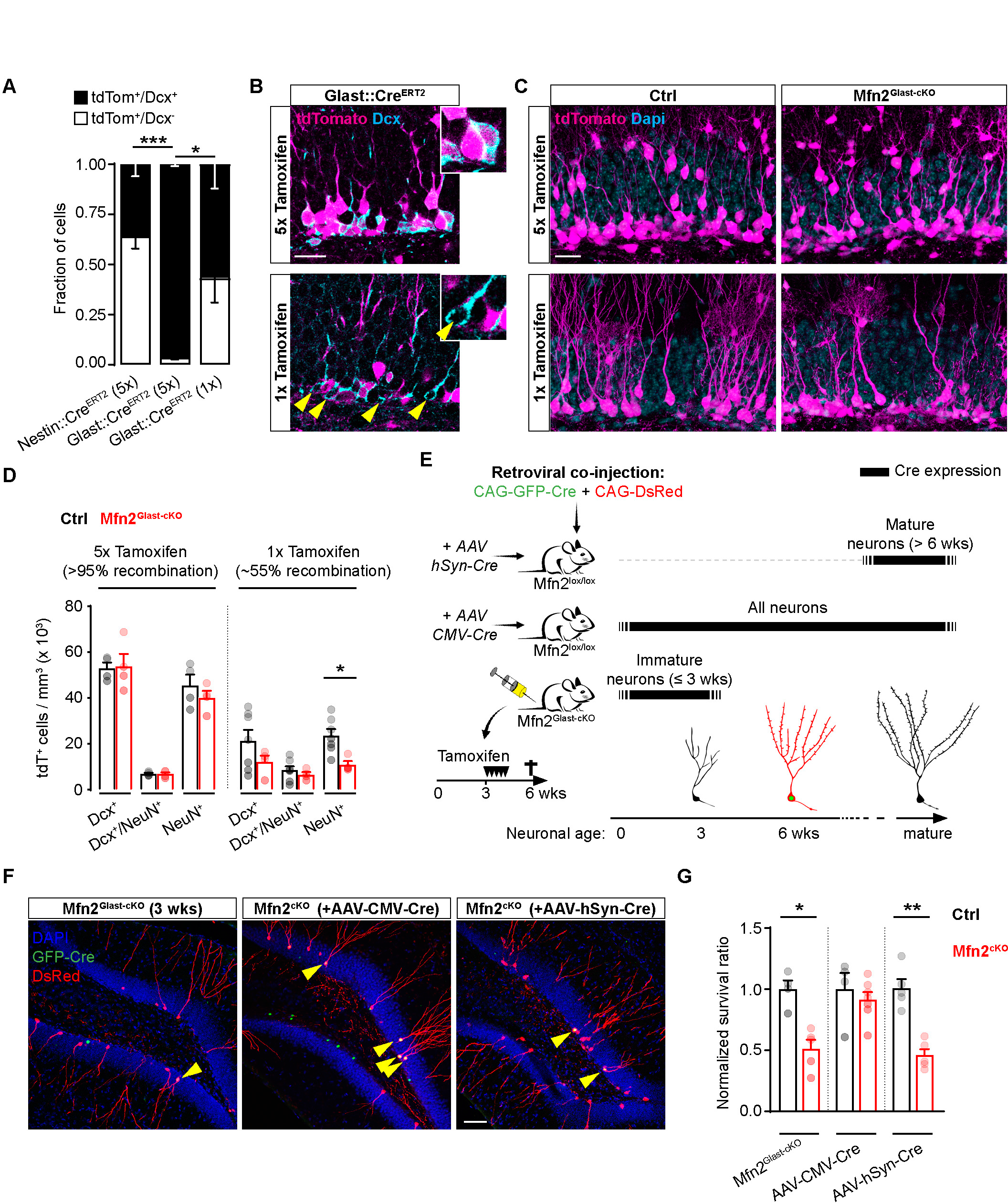
Mitochondrial fusion regulates competition dynamics between adult-born GCs. **(A)** Histogram showing the differences in recombination rate (assessed via tdTomato reporter) in Nestin::Cre^ERT2^ and Glast::Cre^ERT2^ mice following tamoxifen administration once a day for either 1 or 5 days (n= 4-6 mice per condition; two-way Anova followed by Tukey’s multiple comparison test). **(B)** Examples of recombination rate (tdTomato+ cells over Dcx+ immunoreactive immature GCs) in Glast::Cre^ERT2^ mice following either 5 or 1 single dose of tamoxifen. Insets show zooms of Dcx+ cells in the SGZ. Arrowheads point to Dcx+/tdTomato-cells. Bar, 20 µm. **(C)** Examples showing the density of recombined tdTomato+ GCs in the upper DG blade of control and Mfn2^cKO^ mice following either 1 single dose or 5 doses of tamoxifen over 5 consecutive days. Bar, 20 µm. **(D)** Quantification of tdTomato+ GCs expressing Dcx, NeuN or both in control and Mfn2^cKO^ mice following either 1 single dose or 5 doses of tamoxifen over 5 consecutive days (n= 4-7 mice per condition and time point; Mann-Whitney test). **(E)** Experimental settings used for assessing changes in 6-week-old GC survival index following simultaneous ablation of *Mfn2* in the vast majority of GCs older than 6 weeks, younger than 3 weeks or GCs of all ages by means of specific promoter-driven AAV co-injection in Mfn2^lox/lox^ mice or tamoxifen treatment in Mfn2^cKO^ mice. The scheme depicts which age-matched GC populations are targeted by the various approaches. **(F)** Pictures of the DG illustrating examples of co-transduction rates for CAG-DsRed and CAG-GFP-Cre retroviruses according to the combinations depicted in E. Arrowheads point to double-transduced GCs. Bar, 50 µm. **(G)** Quantification of the GC survival index at 6 weeks following retroviral injection according to the combinations depicted in E (n= 4-7 mice per condition; Mann-Whitney test). Data are shown as means ± SEM; *, *P* < 0.05; **, *P* < 0.01; ***, *P* < 0.005; See also Figure S5.

To independently validate these findings, and assess the role of mitochondrial fusion in other potentially competing GC cohorts of age differing (either younger or older) to that 6-week-old GCs, we took advantage of a modified retroviral-based approach to examine the survival index of adult-born GCs lacking *Mfn2* (Figure 5E). Specifically, to evaluate the contribution of older (pre-existing) GCs, we included in the injected viral mix an adeno-associated virus (AAV) encoding for Cre under the human Synapsin (hSyn) promoter, which at the time of injection led to recombination in pre-existing GCs, thus sparing dividing NSPCs and any new GC (Dcx+) that would be generated at later time points (Figures 5E and S5A). Alternatively, we included a ubiquitous (CMV) promoter-driven Cre-encoding AAV to simultaneously ablate *Mfn2* in the vast majority of DG cells. This resulted in widespread neuronal recombination including in Dcx+ GCs of various maturational stages up to 6 weeks after viral injection (Figures 5E and S5A). Lastly, we performed the retroviral injection in Mfn2^Glast-cKO^ mice (devoid of tdTomato reporter to allow the use of DsRed retrovirus), and treated mice with full-dosage tamoxifen 3 weeks later, in order to selectively induce recombination in immature GCs up to and not older than 3 weeks of age at the time of analysis (Figure 5E). Following validation of the used AAV titres to confirm the expected pattern of recombination in the DG with respect to immature Dcx+ GCs (**Figure S5A**) and rule out potential adverse effects on neurogenesis (Johnston et al., 2021) (**Figure S5B and S5C**), we compared the survival ratio of retrovirally birth-dated GCs at 6 weeks of age between these 3 conditions. While restricting Cre-mediated recombination to only younger (Mfn2^Glast-cKO^ mice) or older (hSyn-Cre AAV-infused mice) GCs did not modify the survival index of 6-week-old *Mfn2*-deficient GCs, simultaneous recombination in the vast majority of GCs restored survival to levels comparable to controls (Figure 5F and 5G).

Together, these results strongly suggest that the survival defects in new GCs lacking *Mfn2* are not cell-autonomously regulated, but result from competition dynamics with surrounding wild-type GCs of similar age.

### Disruption of mitochondrial fusion alters experience-dependent circuit incorporation of adult-born GCs

Although the rescuing of GC survival following widespread *Mfn2* deletion in GCs of similar age argues in favour of a role of mitochondrial fusion in regulating neuronal competition dynamics, a key question remains as to how these surviving, but mutated GCs would perform at the circuit level. In other words, how do these *Mfn2*-deficient GCs respond to circuit activity? To answer this question, we first examined how the presynaptic connectome of GCs lacking *Mfn2* would respond to experience. We took advantage of an established rabies virus (RABV)-based, retrograde monosynaptic technique (Wickersham et al., 2007) to map the first-order presynaptic inputs to adult-born GCs in Mfn2^Glast-cKO^ mice exposed to EE for 4 weeks (**Figure S5D**), a paradigm which we have previously shown to elicit a significant remodelling of local and long-range connections to GCs in wild-type animals (Bergami et al., 2015). This method makes use of a G-TVA-DsRed retrovirus for delivering to adult-born GCs a RABV glycoprotein (G) and an avian receptor (TVA) later required by the EnvA-pseudotyped RABV (in this case encoding for GFP) to first infect the TVA-expressing GCs (defined as “starter” cells) and then spread to their first-order presynaptic partners (Deshpande et al., 2013) (Figure 6A). Analysis of GFP-only labelled putative pre-synaptic partners revealed a selective reduction in the connectivity of 6-week-old *Mfn2*-deficient GCs regarding local inputs originating from the DG (most pronounced for local interneurons located in the SGZ/GCL) (Figures 6B and 6E) and the hippocampal CA regions (neurons located throughout hippocampal layers) (Figures 6E and S5E), as well as long-range projections arising from the lateral and medial entorhinal cortex (LEC and MEC) (Figures 6C-6E and S5F), which provide the major excitatory drive to GC dendritic spines in the OML and MML, respectively. Interestingly, no significant changes were detected for other long-range projection neurons (Figure 6E), suggesting that the connectivity alterations of *Mfn2*-deficient GCs in response to experience are input specific.

**Figure 6.**
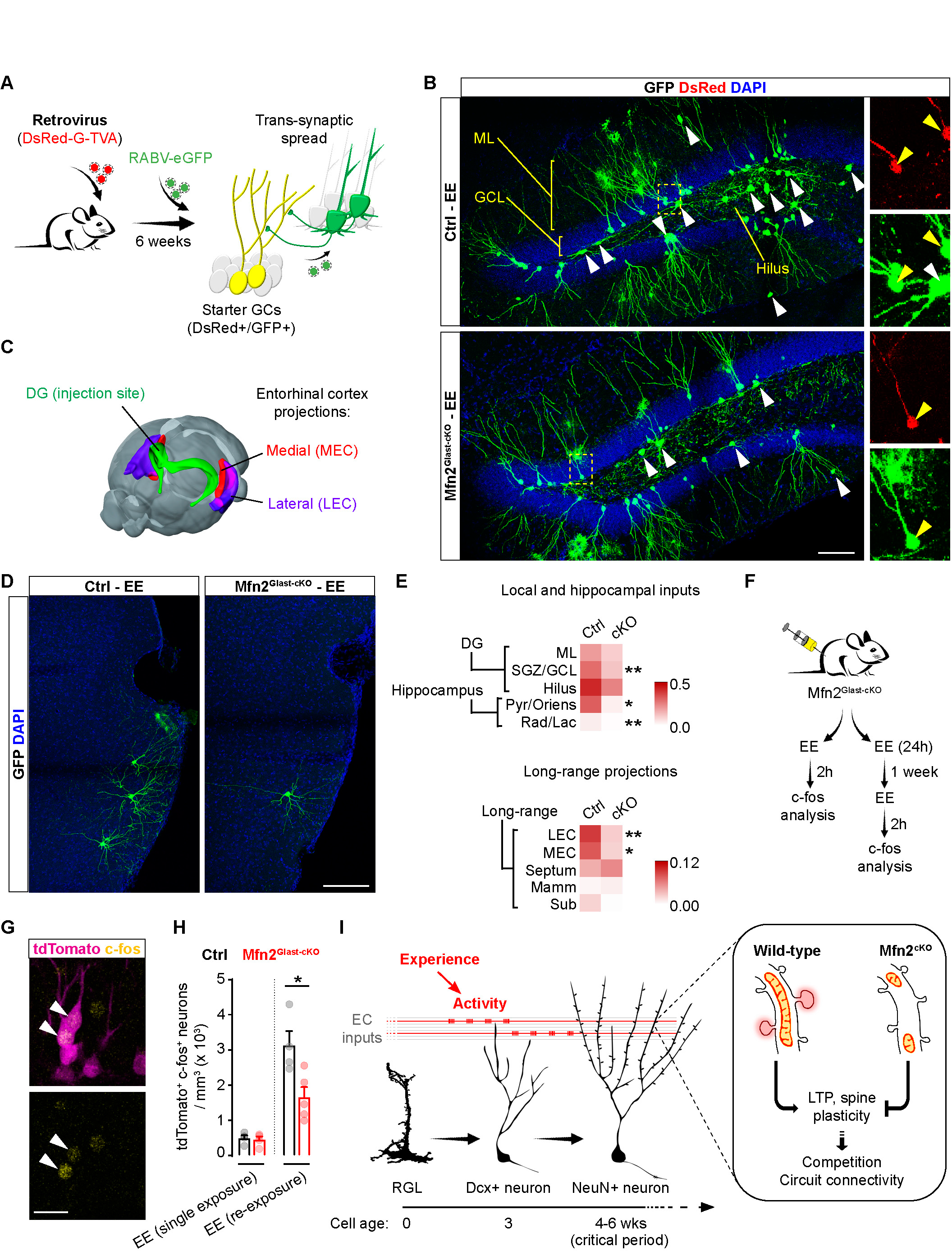
Disruption of mitochondrial fusion alters experience-dependent circuit incorporation of adult-born GCs. **(A)** Experimental design illustrating the RABV-GFP-based trans-synaptic tracing approach to map experience (EE)-dependent connectivity changes of adult-born GCs expressing DsRed, glycoprotein G and TVA receptor. Following initial transduction by DsRed-G-TVA retrovirus, adult-born “starter” GCs are targeted by the EnvA-pseudotyped ΔG RABV encoding for GFP, which then spreads to first-order presynaptic partners. **(B)** Examples of DG sections showing the overall extent of local GFP+ presynaptic connections to adult-born “starter” GCs (DsRed+/GFP+) in control and Mfn2^cKO^ mice following exposure to EE. Panels on the right show zooms of the boxed areas containing starter GCs (yellow arrowheads) and putative presynaptic GFP+ neurons (white arrowhead). Bar, 75 µm. **(C)** Scheme of the mouse brain showing the location of the DG (viral injection site) and regions of the entorhinal cortex (lateral, LEC; and medial, MEC) where long-range excitatory projection neurons are located. **(D)** Examples of traced presynaptic cortical neurons in the LEC of control and Mfn2^cKO^ mice. Bar, 100 µm. **(E)** Heat maps depicting the local and long-range presynaptic connectivity ratio of adult-born GCs quantified according to number and location of presynaptic partners normalized to the number of starter GCs (n= 4-5 female mice per condition; Holm-Sidak multiple t test). **(F)** Experimental paradigm used for assessing adult-born GC activation and memory recall in control and Mfn2^cKO^ mice by c-fos immunoreactivity following either a brief exposure (2h) or a repeated exposure (distanced by 1 week) to the same EE. **(G)** Example showing tdTomato+ recombined adult-born GCs being immunoreactive for c-fos after EE exposure. Bar, 15 µm. **(H)** Quantification of tdTomato+/c-fos+ GCs in control and Mfn2^cKO^ mice following the two paradigms depicted in G (n= 4-5 mice per condition; Mann-Whitney test). **(I)** Graphical summary illustrating the requirement of mitochondrial fusion for adult-born GC synaptic plasticity and circuit connectivity. ML, molecular layer; GCL, granule cell layer. Data are shown as means ± SEM; *, *P* < 0.05; **, *P* < 0.01; See also Figure S5.

Finally, to investigate whether these structural connectivity changes may also mirror functional alterations at the circuit level, we examined network recruitment of adult-born GCs in response to EE by means of immediate-early gene expression (Figure 6F). Analysis of *c-fos* immunoreactivity (Figure 6G), a gene which is induced by synaptic activity and required for synaptic plasticity (Cole et al., 1989; Fleischmann et al., 2003), revealed that *Mfn2*-deficient GCs were less likely than controls to be recruited into behaviourally relevant networks following re-exposure to the same EE, but not following a single brief exposure of 2h (Figures 6F and 6H). This indicates that while wild-type and *Mfn2*-deficient GCs can be similarly activated by a brief exposure to a novel experience, lack of mitochondrial fusion affects GC re-activation by a recently formed memory trace when mice are re-exposed to the same experience.

Taken together, these results support the notion that mitochondrial fusion is required for experience-dependent plasticity of adult-born GC connectivity.

## Discussion

We identified a narrow period during the fourth week of adult-born GC age characterized by enhanced mitochondrial fusion dynamics, which shortly anticipates the critical period of heightened synaptic plasticity between weeks 4 and 6 (Ge et al., 2007). This period of increased mitochondrial remodeling marks a shift from a fragmented and interspersed dendritic network into a more elaborated one, in which mitochondria acquire elongated morphologies, sharply reduce their motility, and stably occupy larger portions of the dendritic shaft, which is consistent with earlier studies performed *in vitro* or in other neuronal populations during early postnatal development (Faits et al., 2016; Rangaraju et al., 2019; Silva et al., 2021). Our findings support the notion that mechanisms controlling the timed acquisition of specific mitochondrial morphologies and metabolic programs contribute to setting the pace of maturation in developing neurons (Iwata et al., 2023) and may in turn facilitate compartment-specific types of synaptic plasticity, which have important implications for cognitive functions associated with adult neurogenesis (Miller and Sahay, 2019). What triggers these changes in mitochondrial fusion during the fourth week of the GC age remains unclear. This may be part of an intrinsic cellular metabolic program meant to remodel mitochondrial morphology as a whole and achieve mitochondrial stabilization throughout the dendritic tree. Alternatively, developmental changes in intrinsic excitability (Marin-Burgin et al., 2012; Mongiat et al., 2009) or the time-matched refinement of locally integrated dendritic signals (Esposito et al., 2005; Ge et al., 2006; Virga et al., 2023) may also play an important role. While conditional deletion of either *Mfn1* or *Mfn2* efficiently prevented this morphological remodeling in dendrites, it did not overly affect the establishment and complexity of the dendritic arbor, neither the formation of spines, whose overall density matched those of control neurons. Yet, ablation of fusion was sufficient to prevent LTP induction and experience-dependent spine plasticity by 6 weeks of the cell age (Figure 6I), while *Mfn2* overexpression was sufficient to boost the formation of mushroom spines in 3-week-old GCs, demonstrating the requirement for fusion in synaptic plasticity of adult-born GCs.

It is somewhat surprising that the overall lineage progression from NSPCs up to the correct morphological development of new mature GCs lacking either *Mfn1* or *Mfn2* was not substantially altered during their first 6 weeks of life. Mitochondrial dynamics, including fusion, have been linked to changes in NSPC and post-mitotic cell fate during embryonic development (Iwata et al., 2020; Khacho et al., 2016). In the adult hippocampus, we found significant changes in NSPC fate and proliferative capacity only following simultaneous deletion of both, *Mfn1* and *Mfn2*, which is consistent with previous work (Khacho et al., 2016), but not in single mutants (despite overt changes in mitochondrial morphology), a finding that corroborates analysis on isolated adult NSPC maintained *in vitro* (Wani et al., 2022). We cannot rule out if the defect in adult NSPC proliferation induced by the combined deletion of *Mfn1* and *Mfn2* may, at least in part, result from mitochondrial dysfunction. This may also potentially explain the distinctive impairment in dendritogenesis of Mfn1/2^cKO^ GCs as compared to single *Mfn* mutants, a phenotype which is reminiscent of that seen in adult-GCs lacking mitochondrial transcription factor A (*Tfam*) (Beckervordersandforth et al., 2017). This therefore suggests that a minimal residual fusion capacity following deletion of either *Mfn1* or *Mfn2* may be sufficient to preserve activity in adult NSPCs.

In newborn GCs, we found that their overall maturation rate (as assessed by marker expression and passive membrane properties) as well as dendrite formation and spinogenesis appeared virtually unaffected by deletion of *Mfn1* or *Mfn2*. Although some differences became manifest when GCs were tested for their excitability (which was reduced in *Mfn2* mutants) as well as frequency and amplitude of spontaneous inputs, functionally the main phenotype was restricted to an impaired synaptic plasticity elicited in response to stimulation, such as LTP induction and mushroom spine formation following EE, which are classic traits of enhanced GC plasticity during their critical period. This is reminiscent of previous work on arcuate nucleus hypothalamic neurons, in which synaptic alterations following deletion of either *Mfn1* or *Mfn2* in mice were revealed only in response to challenge with high-fat diet but not normal chow (Dietrich et al., 2013). While this speaks in favor of potential metabolic rewiring mechanisms that may compensate for basal neuronal activity during protracted lack of mitochondrial fusion (Motori et al., 2020), it also suggests that challenging conditions requiring significant adaptation of synaptic properties may elude such compensatory mechanisms.

In various experimental mouse models, protracted dysregulation of mitochondrial fusion leads to neurodegeneration (Chen et al., 2007; Ishikawa et al., 2019; Lee et al., 2012; Pham et al., 2012), consistent with mutations in the *Mfn2* gene causing the Charcot-Marie-Tooth type 2A disease in humans (Zuchner et al., 2004), a form of peripheral axonal neuropathy. Fate mapping and retroviral birth-dating experiments revealed that deletion of either *Mfn1* or *Mfn2* in only a fraction of adult-born GCs results in a sharp drop of neuronal survival by 6 weeks of age, soon after GCs exited an immature (Dcx+) stage and started acquiring NeuN expression. Yet, clear hallmarks of degeneration only became manifest around 6 months, suggesting the earlier survival defects to be mediated by a “physiological” removal from the neuronal network, and not by a predetermined cell death program. Focusing on *Mfn2*, and consistent with this prediction, we showed that two independent manipulations aimed at simultaneously ablating *Mfn2* in the vast majority newborn GCs of similar age were sufficient to restore neuronal survival. This effect was evident following widespread AAV-mediated knockout of *Mfn2* in adult-born GCs, and in Mfn2^Glast-cKO^ mice by maximizing the extent of tamoxifen-mediated gene recombination in order to target most (>95%) of new GCs. However, no rescue was achieved following restricted Cre-mediated recombination in either fully mature, pre-existing GCs, or more immature newborn GCs that did not yet reach 4 weeks of age, arguing in favor of mitochondrial fusion regulating competition dynamics specifically between adult-born GCs of similar age (4 to 6 weeks). This form of competition differs from previously proposed competition mechanisms between adult-born and pre-existing GCs, which appear to be specifically regulated during the third week of the cell age, namely at the beginning of excitatory innervation, when synaptogenesis peaks and dendritic spines are formed (Kleine Borgmann et al., 2016; McAvoy et al., 2016; Tashiro et al., 2006; Toni et al., 2007). In *Mfn1* or *Mfn2*-deficient GCs, we find that spine formation and overall density are unaffected at all examined time points, suggesting that the initial stages of innervation and spinogenesis proceed as in control GCs. Consistent with this notion, we found no differences in the survival of 3-week-old GCs lacking mitochondrial fusion. Rather, our data point to an effect played by mitochondrial fusion on the plasticity of newly formed spines in mediating a form of competition within GC cohorts of similar age.

The role of *Mfn2* in regulating outer mitochondrial membrane dynamics exceeds mitochondrial fusion. Specifically, *Mfn2* is known to participate in the tethering of mitochondria with the endoplasmic reticulum (ER) to form so-called mitochondria-ER contact sites (or MERCs), at which important metabolic and signaling functions take place (Csordas et al., 2018), including regulation of Ca^2+^ release from the ER, whose disruption can have important consequences for forms of experience-dependent plasticity at dendrites (O’Hare et al., 2022). Therefore, it remains to be addressed whether and to which extent some of the effects described here in adult-born GCs may also be mediated by specific disruption of MERCs. However, the fact that ablation of *Mfn1* alone (which does not participate in MERCs) affects GC survival and the formation of mushroom spines to a similar extent as *Mfn2*, and the fact that GC survival can be rescued by tuning the extent of Cre-mediated recombination in GCs, strongly suggests that these phenotypes are most dependent on changes in mitochondrial fusion rather than MERCs.

## Author contributions

Conceptualization, M.B.; Methodology, S.M.V.K., M.C.M., M.J., H.M.J., G.A.W., F.G., A.S. and M.B.; Investigation, S.M.V.K., M.C.M., H.M.J., G.A.W., F.G., A.S. and M.B.; Formal analysis, S.M.V.K., M.C.M., G.A.W. and M.B.; Writing - Original Draft, M.B.; Writing – Review and Editing, S.M.V.K., M.C.M., H.M.J., G.A.W. and M.B.; Resources, M.J., I.S., D.C.L. and M.B.; Funding Acquisition, M.B.; Supervision, M.B.; Project Administration, M.B.

## Acknowledgments

We thank N.G. Larsson for providing *Mfn1* and *Mfn2* floxed mice as well as *mtYFP* floxed-stop mice. M. Götz and F. Kirchhoff for Glast::Cre^ERT2^ mice. K.K. Conzelmann for providing RABV. J. Matutat, G. Piper, A. Sebbese and the other members of the CECAD in vivo facility for excellent assistance. C. Jüngst and all members of the CECAD imaging facility for assistance with microscopes. J. Göbel and T. Eriksson for help with the initial establishment of Mfn1^GFAP-cKO^ and Mfn2^GFAP-cKO^ lines. V. Sakthivelu for help establishing viral production. This work was supported by the Deutsche Forschungsgemeinschaft (SFB1218 – Grant No. 269925409 – A07, SFB1451 – Grant No. 431549029 – A03 and CECAD EXC 2030 – Grant No. 390661388), European Research Council (ERC-StG-2015, grant number 677844) and Boehringer Ingelheim Stiftung (Exploration Grant) to M.B.

## Author Information

The authors declare no competing financial interests.

## STAR methods

### Lead contact and materials availability

Further information and requests for resources and reagents should be directed to and will be fulfilled by the Lead Contact, Matteo Bergami. All unique/stable reagents generated in this study are available from the Lead Contact without restrictions. There are restrictions to the availability of mice due to MTA.

### Experimental models and subject details

Six to 8-week old C57BL/6N (wild-type) and transgenic mice of mixed genders (unless differently specified) were used in this study. Mice were housed in groups of up to 5 animals per cage supplied with standard pellet food and water *ad libitum* with a 12 h light/dark cycle, while temperature was controlled to 21-22°C. Environmental enrichment (EE) was provided for 4 consecutive weeks by housing mice in larger “hamster” cages (about 76 × 44 cm) equipped with plastic tunnels and other toys, nesting material and running wheels (Bergami et al., 2015). Mice carrying the loxP-flanked genes *Mfn1*^fl/fl^, *Mfn2*^fl/fl^ or both (Lee et al., 2012) were either used as such or crossed with the inducible hGFAP::Cre^ERTM^ (Chow et al., 2008) line and subsequently to the mitochondrial-targeted mtYFP reporter (Sterky et al., 2011). For clonal analysis and population dynamics experiments, *Mfn1*^fl/fl^ or *Mfn2*^fl/fl^ mice were crossed with the Nestin::Cre^ERT2^ line (Lagace et al., 2007) in combination with the inducible tdTomato reporter line Ai14 (Madisen et al., 2010). For all other analyses, *Mfn1*^fl/fl^ or *Mfn2*^fl/fl^ mice were crossed with the Glast::Cre^ERT2^ line (Mori et al., 2006) in combination with the inducible tdTomato reporter line Ai14. All experimental procedures were performed in agreement with the European Union and German guidelines and were approved by the State Government of North Rhine Westphalia.

### Method details

#### Tamoxifen, BrdU and EdU treatments

Mice were intraperitoneally injected with 4-hydroxytamoxifen (40 mg/ml dissolved in 90% corn oil and 10% ethanol) once a day for 1 or 5 consecutive days. For clonal analysis, mice received a single injection of 0.4 mg. The exact time frames of individual experiments are indicated in the text and figures. To examine NSPC proliferation, mice received two i.p. injections of Bromodeoxyuridine (BrdU, 20mg/ml in 0.9% saline) spaced by 4 hours for two consecutive days and sacrificed 24 hours after the last injection. Alternatively, mice were injected i.p. with 5-ethynyl-2’-deoxyuridine (EdU, 20 mg/ml dissolved in 0.9% saline) two times with 2 hours interval and analyzed 2 hours after the last EdU-injection.

#### Stereotactic procedures and viral injections

Mice were anesthetized by intraperitoneal injection of a ketamine/xylazine mixture (100 mg/kg body weight ketamine, 10 mg/kg body weight xylazine), treated subcutaneously with Carprofen (5 mg/kg) and fixed in a stereotactic frame provided with a heating pad. A portion of the skull covering the somatosensory cortex (from Bregma: caudal: −2.0; lateral: 1.5) was thinned with a dental drill to avoid disturbing the underlying vasculature and a small craniotomy sufficient to allow penetration of a glass capillary was performed. For virus injection a finely pulled glass capillary was then inserted through the dura (−1.9 to −1.8 from Bregma) and a total of about 500 nl of volume was slowly infused via a manual syringe (Narishige) in three vertical steps spaced by 50 µm each during a time window of 10-20 minutes. After infusion, the capillary was left in place for few additional minutes to allow complete diffusion of the virus. After capillary removal, the scalp was sutured and mice were placed on a warm heating pad until full recovery. Physical conditions of the animals were monitored daily to improve their wellbeing before euthanizing them.

#### Viral production

The retroviral constructs used in this study were derived from a Moloney Murine Leukemia Virus-based retroviral vector, in which GFP gene expression is driven by the chicken beta-actin promoter (CAG) (Zhao et al., 2006). The CAG-*mTurquois2*, CAG-*mtPA-GFP,* CAG-*GFP-Cre* fusion, CAG-*DsRedExpress2*, CAG-*Cre*-IRES-*DsRedExpress2,* CAG-*Mfn2*-T2A-*GFP-Cre* and the CAG-*Mfn2*-T2A-*GFP* were generated by replacing the GFP coding sequence with the corresponding cDNA of the cytosolic, nuclear or mitochondrial-targeted genes and fluorophores. The retrovirus encoding for *DsRedExpress2*, the RABV *glycoprotein* (*G*) and the *TVA800* (the GPI anchored form of the TVA receptor), designed as CAG-*DsRedExpress2*-2A-*G*-IRES2-*TVA* (i.e., G-TVA retrovirus), as well as the retrovirus encoding for *mtDsRed* with or without *Cre* (CAG-*Cre*-IRES-*mtDsRed*) were described previously (Deshpande et al., 2013; Steib et al., 2014). The chosen retroviral plasmid was used to transfect the helper-free HEK293 gpg cell line using Lipofectamine 2000T, and virus was harvested at 2, 4, and 6 days after transfection, followed by ultracentrifugation. Titers used for experiments were typically in the range of 3 × 10^8^. Construction of the *G* gene-deleted *GFP*-expressing RABV (SADΔG-GFP), virus rescue and pseudotyping have been described before (Deshpande et al., 2013; Wickersham et al., 2007). Recombinant AAV vectors were obtained from Addgene (pENN.AAV.CMVs.Pl.Cre.rBG, cat. 105537-AAV1; and pENN.AAV.hSyn.Cre.WPRE.hGH, cat. 105553-AAV1) and diluted in sterile saline to achieve a concentration of 1 x 10^11^ vg/ml before use.

#### Immunohistochemistry

Mice were anesthetized by intraperitoneal injection of a ketamine/xylazine mixture (130 mg/kg body weight ketamine, 10 mg/kg body weight xylazine), transcardially perfused with 4% PFA in PBS and the brain isolated. Following overnight post-fixation, coronal brain sections (50 µm thick) were prepared using a vibratome (Leica, VT1000 S) and permeabilized in 1% Triton X-100 in PBS for 10 min at RT, followed by brief incubation in 5% BSA and 0.3% Triton X-100 in PBS before overnight immunodetection with primary antibodies diluted in blocking buffer at 4°C on an orbital shaker. The next day, sections were rinsed in PBS 3x 10 min and incubated for 2h at RT with the respective fluorophore-conjugated secondary antibodies diluted in 3% BSA. After washing and nuclear counterstaining with 4’,6-diamidino-2-phenylindole (DAPI, ThermoFisher, 3 µM), sections were mounted on microscopic slides using Aqua Poly/Mount (Polysciences). The following primary antibodies were used: chicken anti-GFP (1:500, Aves Labs, GFP-1020), rabbit anti-RFP (1:500, Rockland, 600401379), rabbit anti-GFAP (1:500, Millipore, ab5804), mouse anti-GFAP (1:500, Millipore, MAB360), mouse anti-c-fos (1:500, Abcam, ab7963), guineapig anti-Dcx (1:1000, Millipore, AB2253), mouse anti-NeuN (1:300, Millipore, MAB377) and mouse anti-Sox2 (1:500, Abcam, ab97959). The following secondary antibodies were used (raised in donkey): Alexa Fluor 488-, Alexa Fluor 546-, Alexa Fluor 647-conjugated secondary antibodies to rabbit, mouse, chicken and rat (1:1000, Jackson ImmunoResearch). For BrdU incorporation analysis, slices were treated for 30 min with 2N HCl at 37 °C and incubated in 0.1 M sodium tetra borate for 15 min at RT before being subjected to incubation with the primary rat anti-BrdU antibody (1:500, Abcam, ab6326) followed by secondary antibody incubation. For EdU incorporation analysis, the Click-iT EdU Imaging kit (Molecular Probes, Invitrogen) was used according to manufacturer’s instructions. For TUNEL assay, the DeadEnd Fluorometric TUNEL System (Promega) was used according to the manufacturer’s instructions.

#### Electrophysiology

Freshly removed mouse brains were placed in ice-cold, carbogen-saturated (5% CO2, 95% O2, pH 7.4) artificial cerebrospinal fluid (aCSF) containing (in mM): 125 NaCl, 2.5 KCl, 1.25 NaH2PO4, 25 NaHCO3, 25 D-Glucose, 0.5 CaCl2 and 3.5 MgCl2, adjusted to pH 7.4 and 310-320 mOsm. 300 μm thick coronal hippocampal slices were prepared with a vibratome (Micron HM650V ThermoScientific, Walldorf, Germany) in ice-cold aCSF. The obtained slices were transferred into a chamber containing aCSF at RT with the following composition (in mM): 125.0 NaCl, 2.5 KCl, 1.25 NaH2PO4, 25.0 NaHCO3, 25.0 D-Glucose, 4.0 CaCl2, 3.5 MgCl2, adjusted to pH 7.4 and 310-320 mOsm. Slices were stored for at least 30 min to allow tissue recovery before starting recordings. All recordings were performed using a microscope stage equipped with a fixed recording chamber and a 40x water-immersion objective (Scientifica). Adult-born GCs were identified by the expression of tdTomato, their morphology, and their anatomical localization in the DG. Patch pipettes with a tip resistance of 4-12 MΩ were made from borosilicate glass capillaries (GB150-10, 0.86 x 2.5 x 100 mm, Science Products, Hofheim, Germany) with a horizontal pipette puller (Model P-1000, Sutter Instruments, Novato, CA, USA). Immediately before recordings, the patch pipette was filled with internal solution containing, in mM: 4.0 KCl, 2.0 NaCl, 0.2 EGTA, 125.0 K-Gluconate, 10.0 HEPES, 4.0 ATP(Mg), 0.5 GTP(Na), 10.0 phosphocreatin, adjusted pH 7.25 and 290 mOsm. Whole-cell patch clamp recordings were performed with an ELC-03XS npi patch clamp amplifier (npi electronic GmbH, Tamm, Germany) that was controlled by the software Signal (version 6.0, Cambridge Electronic, Cambridge, UK). The experiments were recorded with a sampling rate of 12.5 kHz. The signal was filtered with two short-pass Bessel filters that had cut-off frequencies of 1.3 kHz and 10 kHz. Capacitance of the membrane and pipette was compensated using the compensation circuit of the amplifier. All experiments were performed under visual control using the Orca-Flash 4.0 camera (Hammamatsu, Geldern, Germany), controlled by the software Hokawo (version 2.8, Hammamatsu, Geldern, Germany). Criteria to include cells in the analysis were 1) visual confirmation of tdTomato+ signal in the tip of the patch pipette, 2) attachment of the labelled soma to the pipette, 3) and absolute leak current <50 pA at holding potential. Shortly after establishing whole-cell configuration in voltage-clamp mode, the resting membrane potential (E_M_) was measured by applying 0 pA. To determine basic electrophysiological properties different current injection protocols were executed during the recordings. First, spontaneous postsynaptic currents (sPSCs) were recorded for 5 min in voltage-clamp mode with holding potential set to −75 mV. To analyze passive membrane properties, four hyperpolarizing current pulses of −5 pA increments with a duration of 1 s were applied. To analyze action potential properties, depolarizing current pulses of +5 to +10 pA increments with a duration of 2 s were injected. The number of sweeps was individually adjusted for each cell until maximum spike frequency was reached. Finally, an I/V curve was performed by applying 2 s long current injections from −150 pA to +40 pA with +10 pA increments. To perform extracellular stimulation, a monopolar microelectrode, consisting of a chloride-coated silver wire in a patch pipette with a resistance of 1-2 MΩ, was placed under visual control in the medial performant path. To evoke excitatory postsynaptic currents (EPSCs), a baseline protocol was applied that consisted of 100 μs long presynaptic stimulation pulses every 30 s. Stimulation intensity was adjusted individually for each cell to 50% of the maximum EPSC amplitude. Afterwards, EPSCs were recorded for 5 min using 100 μs long presynaptic stimulation pulses every 30 s. This part of the recording served as reference for analysis of short- and long-term plasticity. Short-term synaptic plasticity was evoked by a paired pulse stimulation with 50 ms inter-pulse intervals every 30 s for 5 min. Long-term potentiation (LTP) was evoked with a theta burst stimulation (TBS) paradigm, consisting of four repeated episodes (at 0.1 Hz) of ten stimuli at 100 Hz that were repeated ten times at 5 Hz and paired with a 100 pA postsynaptic current injection. After TBS paradigm, baseline EPSC amplitudes were recorded for at least 40 min with 100 μs long presynaptic stimulation pulses every 30 s. These paradigms were applied for only one cell per slice. Data of electrophysiological recordings were analyzed with Igor Pro (version 7.05.2, WaveMetrics, USA).

#### Ex-vivo live imaging

Isolated brains were placed in ice-cold, carbogen-saturated (5% CO_2_, 95% O_2_, pH 7.4) artificial cerebrospinal fluid (ACSF) containing (in mM): 125 NaCl, 2.5 KCl, 1.25 NaH_2_PO_4_, 25 NaHCO_3_, 25 Glucose, 0.5 CaCl_2_ and 3.5 MgCl_2_ (osmolarity of 310-330). 270-300 µm thick coronal slices were obtained using a vibratome (Micron, HM 650V) and transferred into a pre-incubation chamber maintained at room temperature and containing ACSF supplemented with 1 mM CaCl_2_ and 2 mM MgCl_2_. During imaging, slices were moved in a dedicated imaging chamber and experiments were conducted under continuous ACSF perfusion at a constant temperature of 32-33°C. Imaging in slices was performed using a multiphoton laser-scanning microscope (TCS SP8 MP-OPO, Leica Microsystems) equipped with a Leica 25x objective (NA 0.95, water) and a Ti:Sapphire laser (Chameleon Vision II, Coherent). For mito-PA-GFP experiments, photoactivation of selected dendritic ROIs in individual GCs was carried out by tuning the 2-photon laser to 840 nm (2-5% of laser power for 5 seconds), while time-lapse imaging was performed utilizing GFP excitation (920 nm) and an internal HyD detector (FITC: 500-550 nm). Up to 2 ROIs of similar size per GC were selected in the MML and, after photoactivation, a z-scan (inter-stack interval of 1 μm) the whole volume including ROI and surrounding branches was imaged over the course of at least 20 minutes every 3 minutes. Only GCs located at least 20-30 μm below the slice surface, with a general healthy appearance throughout the recording time (i.e., absence of visibly swollen mitochondria or abnormal appearance of cell bodies) and whose acquisitions displayed only no or a minor spatial drift in xyz during the whole imaging session were included in subsequent analysis. Acquired time points were then merged in a 4D hyperstack in ImageJ and the resulting 3D volumes registered utilizing the “Correct 3D drift” plugin in ImageJ. Quantification of fusion events was performed manually by inspecting the registered time-lapse acquisitions including and surrounding the photoactivated ROIs. Fusion events were identified by the abrupt decrease in GFP intensity in directly photoactivated mitochondria due to GFP dilution into the newly appearing (fusing) mitochondria that had not been initially photoactivated. Photoactivated mitochondria that simply moved away or though the ROIs and did not satisfy these fusion parameters were considered as motility events.

#### Image analysis and quantification

Images were acquired utilizing SP8 Confocal microscopes (Leica Microsystems) equipped with 10x (air objective, NA 0.3), 20x (NA 0.75), 40x (NA 1.3), 63x (NA 1.2) and 100x (NA 1.4) immersion objectives, white light laser and multiple HyD detectors. For calculation of the survival index following retroviral birth-dating transduction, the whole hippocampus was sectioned (50 to 70μm thick coronal sections) and single and double positive transduced GCs for the utilized virally-encoded fluorophore were manually quantified per mouse. For quantification of cell proliferation and GC densities, 4 to 6 coronal brain sections per brain corresponding to similar anatomical locations across mice were used. All acquired z-stack images (LAS software) were converted into TIFF files and analysis was performed off-line using the ImageJ software (National Institute of Health, Bethesda, United States). Cells counting was performed manually by using the Cell Counter plugin and by normalizing the number of marker+ cells over the volume of the GCL in the DG (measured area multiplied by the inter-stack interval). The total number of RGL in the DG was obtained by examining their unique morphological features, which makes them well distinguishable from other cell types (short radial process spanning the granule cell layer, small body in the SGZ), and positivity for the marker GFAP or Sox2. Likewise, the number of GCs was assessed by unique morphological features (round cell body and a visible dendritic arbor) as well as by the positivity for reporter gene (i.e., tdTomato), immature (Dcx) or more mature markers (NeuN). For clonal analysis, the whole hippocampus was investigated through serial brain sections (thickness 75µm), which were first screened for TdTomato+ cells with a 20x oil objective and then z-stack acquisition of regions containing positive cells taken with a 20x and a 63x oil objectives. For quantification, regions including the molecular layer (ML), granular cell layer (GCL), sub granular zone (SGZ) and Hilus bordering the SGZ were considered. A radius of 150 μm from the RGL cell within the clone was used to determine the spatial limits of the clone itself (Bonaguidi et al., 2011). Clones were categorized according to the presence or absence of RGL and by the composition of other cell types (neurons and astrocytes). After initial validation by marker expression of possible tdTomato+ cell types (see above), cell identity in individual clones was determined based on morphology: RGL with a triangular-shaped soma located in the SGZ, a short radial process branching in the inner molecular layer and absence of any axon; neurons for having round cell body located in in the GCL and clearly visible apical dendritic processes; and protoplasmic astrocytes, for having a bushy morphology irrespective of their locations. For imaging of mitochondria in RGL *in vivo*, mice bearing the mtYFP reporter were utilized, and z-stack acquisitions of individual RGL in the upper blade of the DG (identified by location of their soma in the SGZ and the presence of a distinct GFAP+ radial process) taken with a 63x oil objective utilizing an inter-stack interval of 0.3 μm. For trans-synaptic tracing analysis, the whole brain was sectioned (70mm thick coronal sections) and quantifications were performed by counting the number of double-transduced (GFP+ and DsRed+) as well as RABV-only transduced cells (GFP+) per mouse. The connectivity ratio (or converge index) for each anatomical location (layers of the DG and hippocampus, as well as long-range projection areas) was calculated by normalizing the absolute number of RABV-only transduced presynaptic partners onto the number of double-transduced starter GCs to take into account for changes in the number of starter neurons. For 3D-reconstruction of dendritic and mitochondrial volumes, z-stack of retrovirally labeled newborn GCs were acquired at 1024×1024 pixel resolution with a magnification of 63x, a digital zoom of 3 and a 0.3 µm inter-stack interval. For analysis, the recorded confocal stacks were imported into Imaris software (version 8.3.1, Bitplane) and rendered in 3D to generate volume masks for both the dendritic and the mitochondrial volume, following by calculation of mitochondrial sphericity index and occupancy ratios. For dendritic tree reconstruction and Sholl analysis, the NeuronJ plugin for ImageJ was used on confocal images to manually trace the dendritic morphology of selected GCs and measure their overall dendritic length as well as branching points. The reconstructed neuron shapes were converted into binary images in ImageJ and used for Sholl analysis (Neuroanatomy function in ImageJ, Legacy Sholl analysis with a radius step size of 5 microns). For spine density analysis of adult-born GCs, serial z-stacks (0.3μm inter-stack interval) of medial parts of the dendritic tree within the superior blade of the DG were acquired with a 63x objective (PL Apo 63x/1.2 water DIC) with a digital zoom of 3x. Continuous dendritic segments were visualized in a maximum projection, and ImageJ was used to quantify linear spine density and to estimate the individual cross-sectional area of large spines (mushroom spines were classified for having an area equal or higher to 0.4 μm^2^).

#### Correlative light and electron microscopy

Sections were stained for nuclei with 4’,6-diamidino-2-phenylindole (DAPI, ThermoFisher, 3 µM) and placed into imaging dishes with glass bottom (ibidi, Graefelfing, Germany) filled with PBS. Sections were prevented from floating using brain slice anchor (Warner Instruments) and z-stacks of the region of interest in DG were acquired using a SP8 confocal microscope (Leica). After confocal imaging, sections were prepared for transmission electron microscopy using standard protocols. Briefly, post-fixation was applied using 1% Osmiumtetroxid (Science Services, München, Germany) and 1.5% Potassium hexacyanoferrat (Merck, Darmstadt, Germany) for 30 min at 4°C. After 3×5min washes with ddH_2_O, samples were dehydrated using ascending ethanol series (50%, 70%, 90%, 100%) for 10 min each. Infiltration was carried out with a mixture of 50% Epon/ethanol for 1h, 70% Epon/ethanol for 2h and overnight with pure Epon (Merck, Darmstadt, Germany). After fresh Epon for 4h, sections were mounted onto empty polymerized epon blocks and covered with Aclar foil to provide a flat surface. After 48h hardening at 60°C, aclar foil was removed and samples were trimmed to the region of interest which was previously acquired by confocal microscopy using a diamond 90° trimming tool (Diatome, Biel, Switzerland). For orientation, stereotypic shapes of the hippocampus including the granule cell and molecular layers were used and matched to measurements obtained from confocal images of the same region. When the desired z-position was reached, serial sections (50nm) were cut using an UC6 ultramicrotome (Leica, Wetzlar, Germany) and collected onto pioloform (Plano, Wetzlar, Germany) coated slot grids. Post-staining was performed with 1.5 % uranyl acetate (Agar Scientific, Stansted, United Kingdom) for 15 min and Reynolds lead citrate (Roth, Karlsruhe, Germany) solution for 3 min. Electron micrographs were acquired using a JEM-2100 Plus Transmission Electron Microscope (JEOL, Tokio, Japan) operating at 80kV equipped with a OneView 4K camera (Gatan, Pleasanton, USA). Registration of images obtained by light (confocal) and electron microscopy was done using nuclei of blood vessels and nuclei (with nucleoli) of neurons as fiducials with the plugin EC-CLEM from the software ICY (Paul-Gilloteaux et al., 2017).

### Quantification and statistical analysis

Data are represented as means ± SE. Graphical illustrations and significance were obtained with GraphPad Prism 7 (GraphPad). Significance was calculated as described in each figure legend.

## Legend to Supplemental Figures

**Figure S1.**
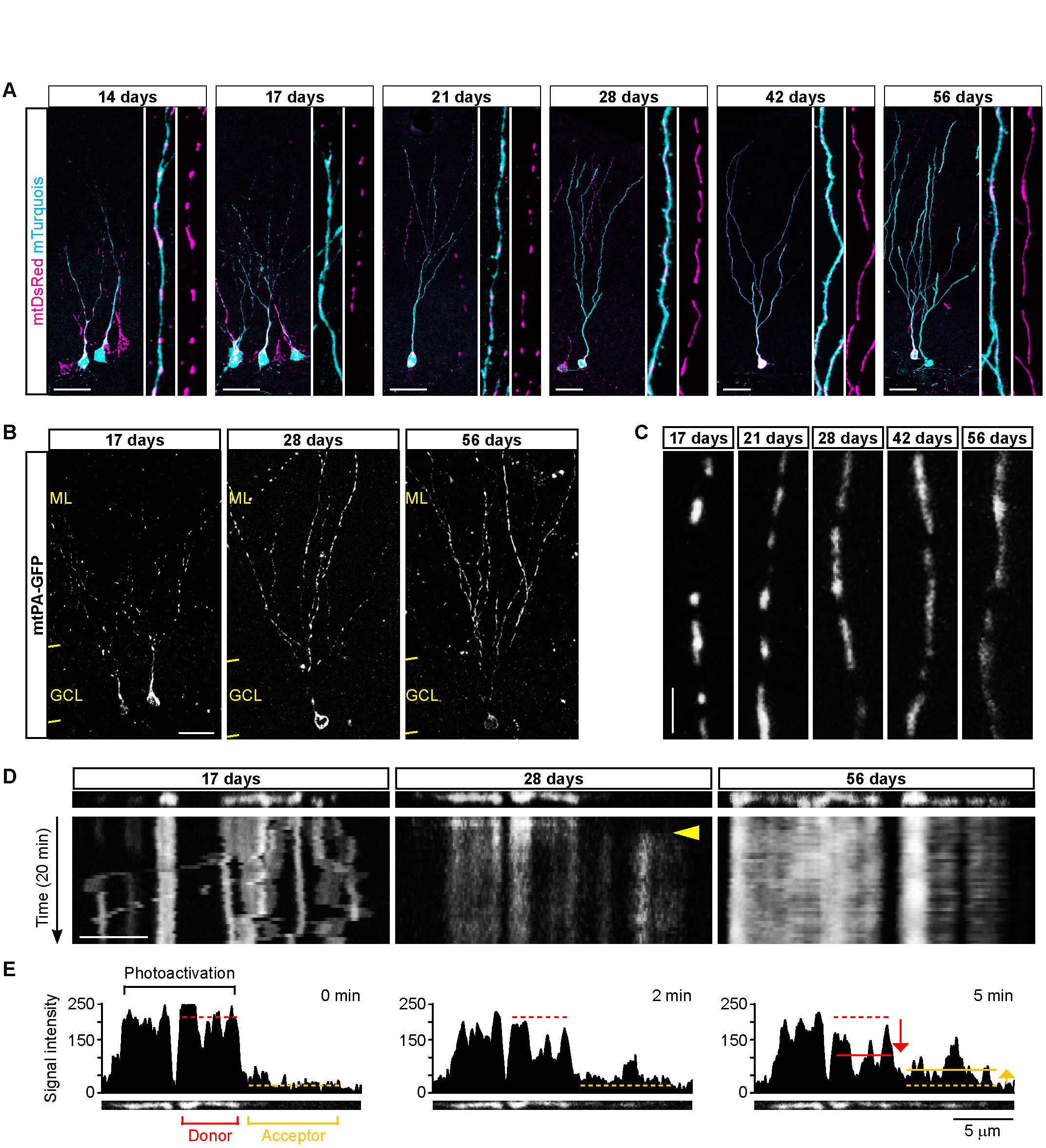
Related to Figure 1. Mitochondrial morphology and dynamics in adult-born GCs. **(A)** Examples of GCs at the indicated ages after retroviral transduction showing changes in the dendritic mitochondrial network (labelled via mtDsRed) with respect to the dendritic tree (labelled with cytosolic mTurquois). Side panels depict zooms of the dendrite. Bar, 30 µm. **(B)** Representative images of adult-born GCs virally transduced with mtPA-GFP and acquired by live imaging via 2PLSM at the indicated ages. **(C)** Zoom of GC dendrites expressing mtPA-GFP in the MML of the DG acquired by live imaging via 2PLSM at the indicated ages. Bar, 5 µm. **(D)** Kymographs of GC dendrites expressing mtPA-GFP in the MML of the DG acquired by live imaging (one scan every 3 minutes for 20 minutes) via 2PLSM at the indicated ages. Mitochondrial motility is evident as diagonal traces, while fusion events (yellow arrowheads) appear as sudden changes in signal intensity of “donor” (photoactivated signal reduction) and nearby non-photoactivated “acceptor” mitochondria (signal appearing). For comparison, a stable cluster of photoactivated mitochondria undergoing no visible dynamics is shown at 56 days. Bar, 5 µm. **(E)** Histogram analysis of signal intensity in the photoactivated ROI and surrounding region shown in D at 28 days after viral transduction. Histograms depict 3 time points before, during and after a fusion event. The delta in GFP signal reduction and increase of donor and acceptor mitochondria, respectively, are shown.

**Figure S2.**
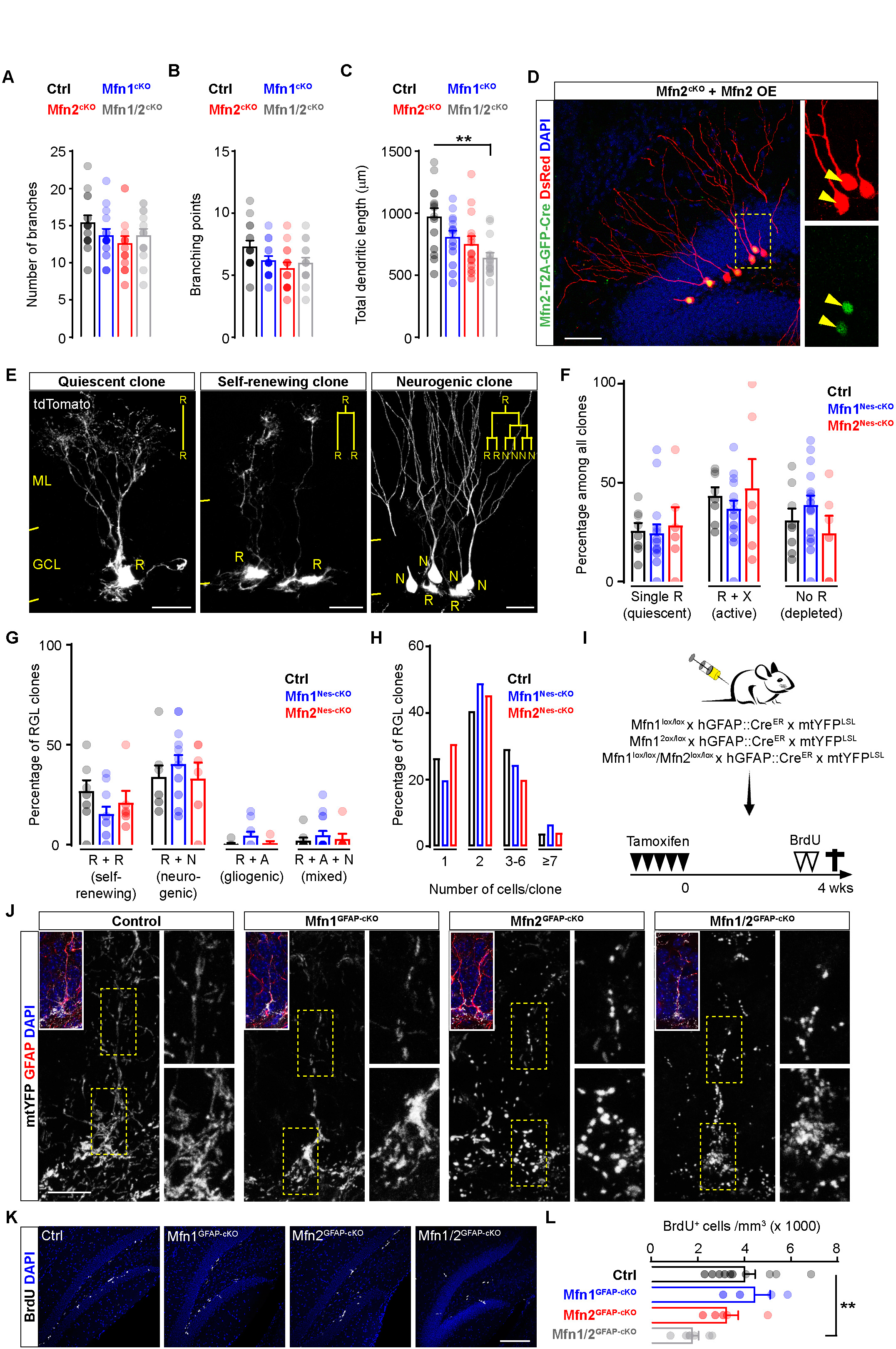
Related to Figures 2 and 3. Disruption of *Mfn1* or *Mfn2* in adult neurogenesis. **(A-C)** Quantification of number of branches, branching points and total dendritic length in transduced Mfn1^cKO^, Mfn2^cKO^ and Mfn1/2^cKO^ GCs (n= 15 neurons obtained from 3-5 mice per genotype; Kolmogorov-Smirnov test). Data are shown as means ± SEM; *, *P* < 0.05; **, *P* < 0.01; ***, *P* < 0.005; Kruskal-Wallis followed by Dunn’s multiple comparison test). **(D)** Example of adult-born GCs co-transduced with CAG-DsRedExpress2 and CAG-Mfn2-T2A-GFP-Cre retroviruses showing lack of abnormal morphological features. Side panels depict a zoom of the boxed area containing double-transduced GCs (yellow arrowheads). Bar, 50 µm. **(E)** Examples of individual clones at 1 month after tamoxifen administration. Putative lineage outcomes of the initially recombined radial-glia like NSPC (R) is shown. Bars, 20 µm. **(F)** Quantification of clones containing a single R (quiescent), R+X (active and self-renewing) and no R (depleted) for the indicated genotypes (n=8 mice per group, Two-way Anova followed by Tukey’s multiple comparison test). **(G)** Proportion of R-containing clones classified according to their cellular composition (n=8 mice per group, Two-way Anova followed by Tukey’s multiple comparison test) (N, neuron; A, astrocyte). **(H)** Quantification of average clone size per genotype (n= 8 mice per genotype). **(I)** Experimental paradigms used for examining mitochondrial morphology and proliferative activity of NSPCs in tamoxifen-inducible Mfn1^GFAP-cKO^ and Mfn2^GFAP-cKO^ mice. **(J)** Examples of radial glia-like mtYFP+/GFAP+ NSPCs (see inset) in the DG of the indicated genotypes. Arrowheads points to the cell soma. Side panels show zooms of the boxed areas in the cell soma and along the main radial process. Bar, 15 µm. **(K)** Examples of BrdU labelling in the DG of the indicated genotypes. Bar, 120 µm. **(L)** Quantification of NSPC proliferation for the indicated genotypes (n=4-11 mice per group; Kruskal-Wallis followed by Dunn’s multiple comparison test). Data are shown as means ± SEM; **, *P* < 0.01.

**Figure S3.**
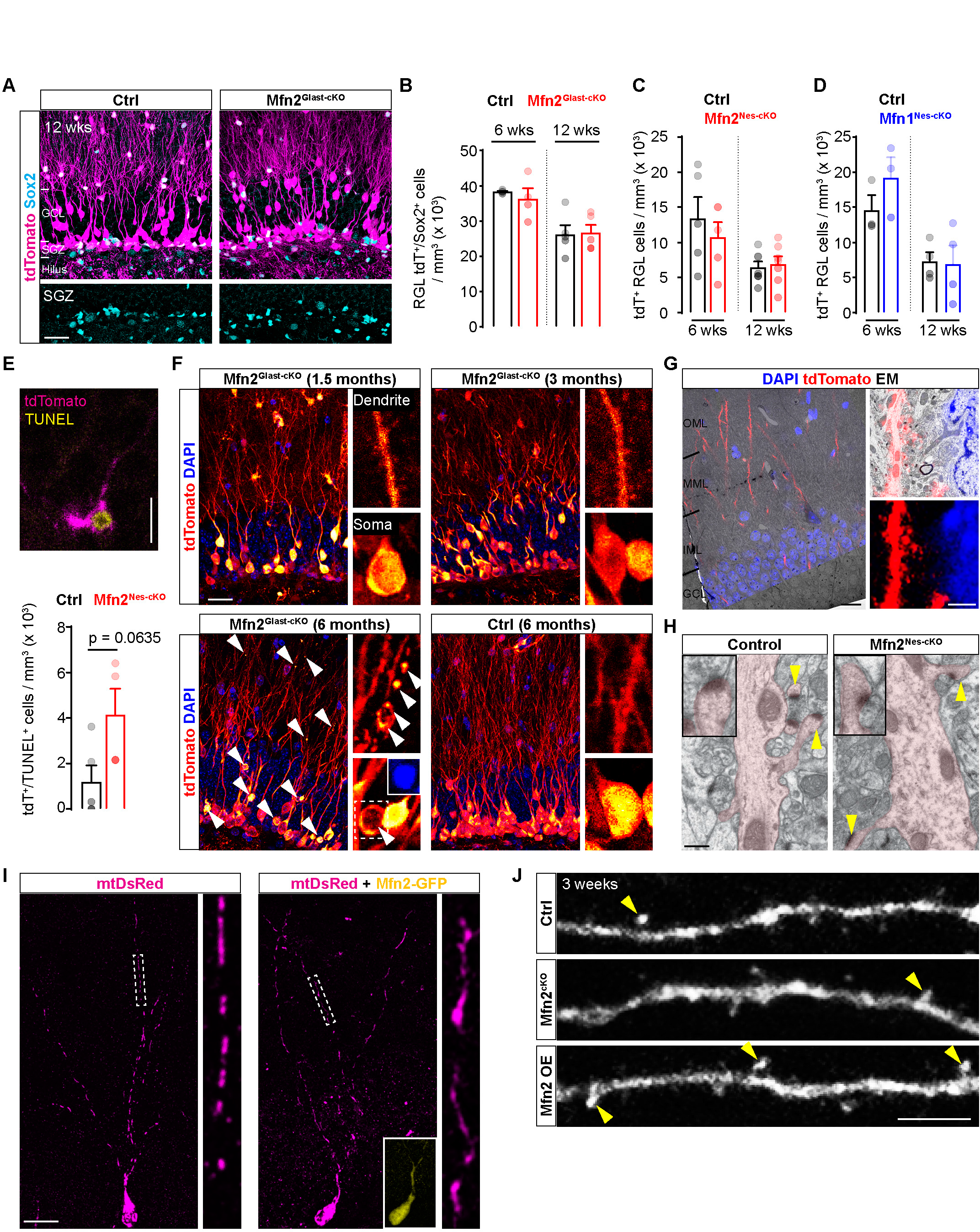
Related to Figures 3 and 4. Effects of *Mfn2* manipulation on RGL cells numbers and adult-born GC survival. **(A)** Examples of the upper DG blade in control and Mfn2^Glast-cKO^ mice showing the density of tdTomato+ and Sox2+ cells at 6 weeks after tamoxifen administration. Bar, 25 µm. **(B)** Quantification of tdTomato+ RGL cells co-expressing the NSPC marker Sox2 in the SGZ of the DG of control and Mfn2^Glast-cKO^ mice at 6 and 12 weeks (n=4-5 mice per group, Mann-Whitney test). **(C-D)** Quantification of morphologically identified tdTomato+ RGL cells in the SGZ of the DG of control, Mfn2^Nes-cKO^ (n=5-7 mice per group) and Mfn1^Nes-cKO^ mice at 6 and 12 weeks (n=3-4 mice per group, Mann-Whitney test). **(E)** Example of a tdTomato+/TUNEL+ adult-born GC in a Mfn2^Nes-cKO^ mouse. Bar, 20 µm. The lower histogram depicts the density in tdTomato+/TUNEL+ cells in control and Mfn2^Nes-cKO^ mice. (n=4-5 mice per group, Mann-Whitney test). **(F)** Examples of control and Mfn2^Glast-cKO^ mice over the course of 6 months after tamoxifen-induced recombination showing the late appearance of apoptotic GCs (white arrowheads) in absence of *Mfn2*. Side panels depict zooms of the boxed areas for GC soma and dendrite. Bar, 25 µm. **(G)** CLEM (low magnification) of adult-born GC dendrites expressing endogenous tdTomato in Mfn2^Nes-cKO^ mice. DAPI staining was used as reference for registration of the fluorescence signal with the electron micrograph. Bar, 10 µm. **(H)** Examples of CLEM-identified GC dendrites (highlighted by a manual masking) in control and Mfn2^Nes-cKO^ mice. Insets report on zooms of spines in each example (yellow arrowheads). Bar, 0.2 µm. **(I)** Examples of mtDsRed-expressing newborn GCs at 3 weeks after transduction in absence or presence of co-transduction with CAG-Mfn2-T2A-GFP. Side panels show zooms of the boxd regions in the dendritic tree. Bar, 20 µm. **(J)** Examples of dendritic segments in 3-week-old newborn GCs of the indicated conditions showing the abundance of mushroom spines (arrowheads). Bar, 5 µm. Data are shown as means ± SEM.

**Figure S4.**
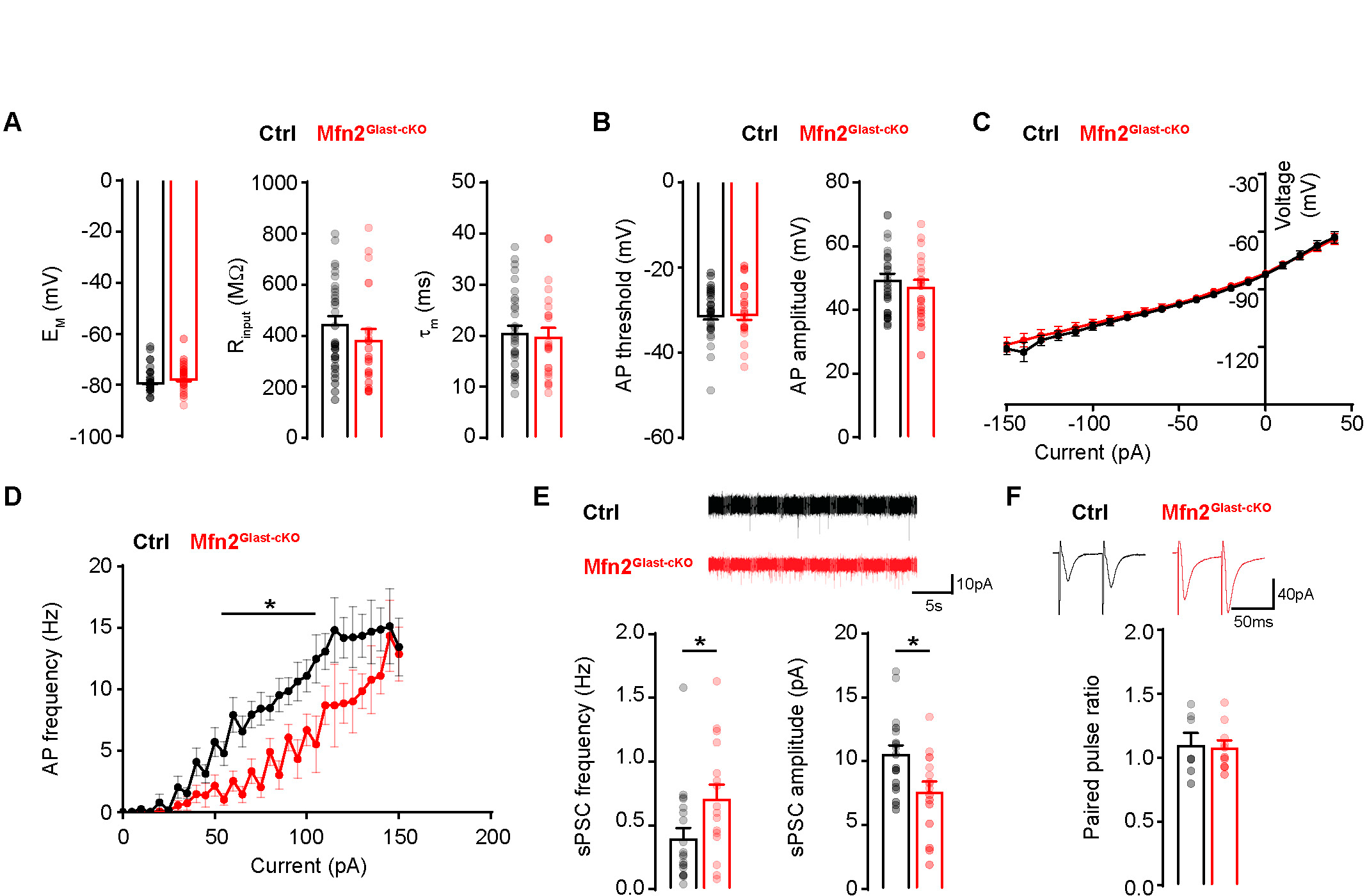
Related to Figure 4. Electrophysiological properties of adult-born GCs lacking *Mfn2*. **(A)** Comparison of passive membrane properties in control and Mfn2^Glast-cKO^ GCs including mean resting membrane potential (E_M_) (n=27-37 cells per group), input resistance (R_input_) (n=22-32 cells per group) and membrane time constant (τ_m_) (n=24-32 cells per group; Mann-Whitney test). **(B)** Comparison of threshold to action potential (AP) and mean AP amplitude in control and Mfn2^Glast-cKO^ GCs following depolarizing current injection (n=21-29 cells per group; Mann-Whitney test). **(C)** Current-voltage relationship (IV curve) in control and Mfn2^Glast-cKO^ GCs (n=14-19 cells per group). **(D)** Average AP frequency in control and Mfn2^Glast-cKO^ GCs following sequential depolarizing current injection (n=22-30 cells per group; Two-way Anova followed by Bonferroni’s multiple comparisons test). **(E)** Quantification of frequency and amplitude of spontaneous postsynaptic currents (sPSCs). Upper traces show representative recordings of sPSCs in control and Mfn2^Glast-cKO^ GCs (n=16-20 cells per group; Mann-Whitney test). **(F)** Quantification of paired-pulse ratio following stimulation of the MML. Upper traces show representative responses induced by two sequential stimuli (50 ms apart) in control and Mfn2^Glast-cKO^ GCs (n=6-7 cells per group; Mann-Whitney test). Data are shown as means ± SEM; *, *P* < 0.05.

**Figure S5.**
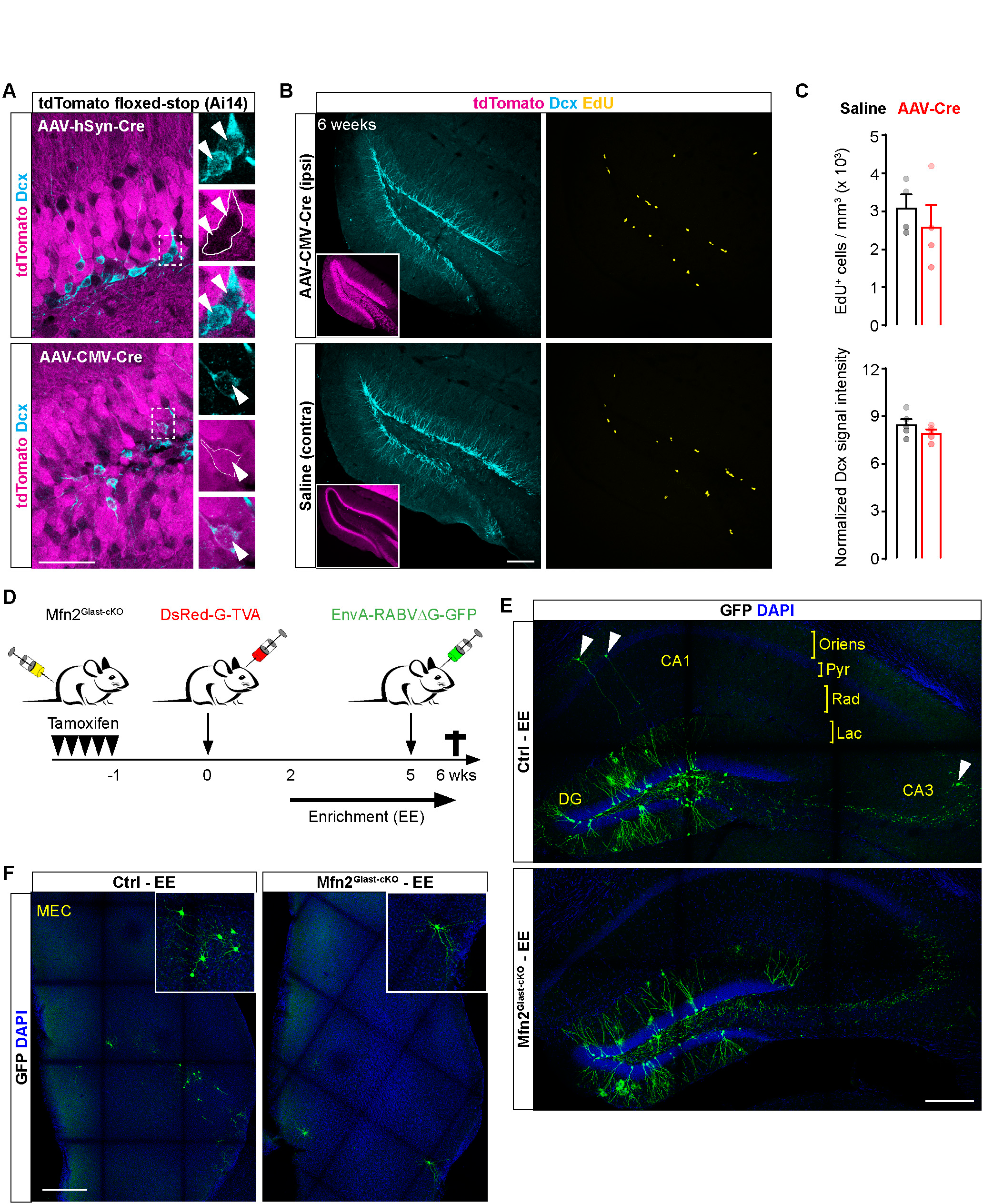
Related to Figures 5 and 6. Validation of AAV-based, Cre-mediated recombination in targeting mature and young GCs in the DG. **(A)** Single confocal stack examples showing the GCL of the DG in tdTomato floxed-stop (Ai14) mice 6 weeks after infusion of the AAV-hSyn-Cre (upper panels) or AAV-CMV-Cre (lower panels). Analysis of Cre-mediated recombination leading to tdTomato expression is shown in zooms with respect to (Dcx+) newborn GCs (lacking or expressing tdTomato in AAV-hSyn-Cre and AAV-CMV-Cre injected mice, respectively). Bar, 50 µm. **(B)** Representative examples of ipsi- and contralateral hemispheres in AAV-CMV-Cre injected tdTomato floxed-stop mice showing the immunoreactivity for Dcx+ immature GCs and proliferating NSPCs following EdU injection the last 2 days before sacrifice. Bar, 70 µm. **(C)** Quantification of EdU+ cells and Dcx immunoreactivity in AAV-CMV-Cre and saline injected hemispheres as shown in B (n=5 mice; Mann-Whitney test). **(D)** Scheme showing the experimental paradigm used to examine changes in the presynaptic connectivity of wild-type and *Mfn2*-deficient adult-born GCs via RABV-based trans-synaptic tracing and during exposure to environmental enrichment (EE). **(E)** Examples of hippocampal sections corresponding to the pictures shown in Figure 6B showing the overall extent of GFP+ presynaptic connections (arrowheads) to adult-born “starter” GCs (DsRed+/GFP+) distributed across the hippocampal layers in control and Mfn2^cKO^ mice following exposure to EE. Bar, 150 µm. **(F)** Examples of traced presynaptic cortical neurons in the MEC of control and Mfn2^cKO^ mice. Bar, 100 µm. Pyr, stratum pyramidale; Rad, stratum radiatum; Lac, stratum lacunosum moleculare. Data are shown as means ± SEM.

